# Co-expressed subunits of dual genetic origin define a conserved supercomplex mediating essential protein import into chloroplasts

**DOI:** 10.1101/2020.07.04.188128

**Authors:** Silvia Ramundo, Yukari Asakura, Patrice A. Salomé, Daniela Strenkert, Morgane Boone, Luke C. M. Mackinder, Kazuaki Takafuji, Emine Dinc, Michèle Rahire, Michèle Crèvecoeur, Leonardo Magneschi, Olivier Schaad, Michael Hippler, Martin C. Jonikas, Sabeeha Merchant, Masato Nakai, Jean-David Rochaix, Peter Walter

**Affiliations:** Department of Biochemistry and Biophysics, University of California at San Francisco, San Francisco, USA; Howard Hughes Medical Institute; Laboratory of Organelle Biology, Institute for Protein Research, Osaka University, Osaka, Japan; Department of Chemistry and Biochemistry, UCLA, Los Angeles, USA; Present address: QB3, University of California, Berkeley, USA; Department of Biology, University of York, UK; Graduate School of Medicine, Osaka University, Osaka, Japan; Departments of Molecular Biology and Plant Biology, University of Geneva, Geneva, Switzerland; Institute of Plant Biology and Biotechnology, University of Münster, Münster, Germany; Department of Biochemistry, University of Geneva, Geneva, Switzerland; Institute of Plant Science and Resources, Okayama University, Kurashiki, Japan; Department of Molecular Biology, Princeton University, USA

## Abstract

In photosynthetic eukaryotes, thousands of proteins are translated in the cytosol and imported into the chloroplast through the concerted action of two translocons — termed TOC and TIC — located in the outer and inner membranes of the chloroplast envelope, respectively. The degree to which the molecular composition of the TOC and TIC complexes is conserved over phylogenetic distances has remained controversial. Here, we combine transcriptomic, biochemical, and genetic tools in the green alga Chlamydomonas (*Chlamydomonas reinhardtii*) to demonstrate that, despite a lack of evident sequence conservation for some of its components, the algal TIC complex mirrors the molecular composition of a TIC complex from *Arabidopsis thaliana.* The Chlamydomonas TIC complex contains three nuclear-encoded subunits, Tic20, Tic56 and Tic100, and one chloroplast-encoded subunit, Tic214, and interacts with the TOC complex, as well as with several uncharacterized proteins to form a stable supercomplex (TicToc), indicating that protein import across both envelope membranes is mechanistically coupled. Expression of the nuclear and chloroplast genes encoding both known and the here newly identified TicToc components is highly coordinated, suggesting that a mechanism for regulating its biogenesis across compartmental boundaries must exist. Conditional repression of Tic214, the only chloroplast-encoded subunit in the TicToc complex, impairs the import of chloroplast proteins with essential roles in chloroplast ribosome biogenesis and protein folding and induces a pleiotropic stress response, including several proteins involved in the chloroplast unfolded protein response. These findings underscore the functional importance of the TicToc supercomplex in maintaining chloroplast proteostasis.

## Introduction

Chloroplasts are vital organelles in eukaryotic photosynthetic organisms. Akin to mitochondria, they are thought to have arisen from an endosymbiotic event, in which a cyanobacterial ancestor was engulfed by a eukaryotic cell (*1*). Over time, most cyanobacterial genes were transferred to the nuclear genome of the host, which nowadays supplies the organelle with the majority of its resident proteins from the cytosol. An essential step in the establishment of chloroplasts was the evolution of the translocons that select chloroplast precursor proteins synthesized in the cytosol and import them through protein-conducting channels into the chloroplast. The translocons are multiprotein complexes, located in the outer and inner chloroplast envelope membranes. Receptor proteins intrinsic to the translocons recognize a signal-peptide (for chloroplasts and mitochondria also called transit-peptide) at the N-terminus of chloroplast precursor proteins that is cleaved off and quickly degraded upon import [original findings: (*2–12*); recent reviews: (*13–19*)]. Based on the membrane in which the translocons reside, they are referred to as translocon of the outer and inner chloroplast membrane (TOC and TIC), respectively (*20*).

To date, the vast majority of the translocon components have been identified and characterized through biochemical studies in green peas (*Pisum sativum*) (*8, 21–30*) and genetic studies in Arabidopsis (*Arabidopsis thaliana*) (*11, 29, 31-37*). Their evolutionary conservation in other photosynthetic eukaryotes has been inferred from phylogenetic sequence alignments (*15*). The molecular composition of the TOC complex is relatively well understood. It consists of three core subunits, namely the pre-protein receptors Toc159 and Toc34, and the protein-conducting channel, Toc75 (*11, 38–41*). By contrast, the molecular identity and function of the subunits in the TIC complex is still under debate. Earlier studies performed on pea chloroplasts revealed several of its subunits, including Tic110, Tic62, Tic55, Tic40, Tic32, Tic22, and Tic20 (*21, 24–28*). However, there is no consensus as to which components form the protein-conducting channel: while multiple studies demonstrated a channel activity for Tic20 *in vitro* (*42–45*), it is still debated whether Tic110 is also directly involved in forming a channel (*46–49*). The scenario has been further complicated by the recent isolation of a novel large TIC complex in Arabidopsis, termed the “1-MDa complex” (*42, 44*), that contains Tic20 in association with a different set of proteins, namely Tic100, Tic56 and Tic214. Tic214 is unique in that it is the only known translocon component encoded by the chloroplast genome (*44*). The 1-MDa TIC complex associates with translocating pre-proteins, displays pre-protein-dependent channel activity (*44*), and functionally and physically cooperates with the Ycf2/Ftshi complex, a recently-identified ATP-driven import motor (*50*). However, two recent studies reported that chloroplast protein import is only partially impaired in Arabidopsis mutants lacking Tic56 (*51*), as well as in Arabidopsis plants in which Tic214 was down-regulated to undetectable levels upon treatment with spectinomycin, a drug that selectively inhibits chloroplast and mitochondrial translation (*52*). Furthermore, no clear orthologue for Tic214 has been identified in grasses, glaucophytes, red algae, or some dicots (*52–55*). Hence, published work raises skepticism whether the 1-MDa TIC complex in general, and Tic214 in particular, are functionally important during chloroplast protein import.

Here, we address these questions using multipronged and unbiased approaches. Using co-immunoprecipitation analyses and co-expression studies, we demonstrate that in Chlamydomonas, all the subunits of the Arabidopsis 1-MDa TIC complex are conserved and are part of a supercomplex that contains all known TOC subunits, as well as several novel subunits. Furthermore, we show that the supercomplex is functional *in vitro* and *in vivo*, and is vital to maintain chloroplast proteostasis.

## Results

### The molecular architecture of plant TOC and TIC is conserved in algae

To establish a comprehensive inventory of TIC and TOC components in Chlamydomonas, we first used the Arabidopsis TOC and TIC protein sequences and queried the Chlamydomonas proteome using the Basic Local Alignment Search Tool (BLAST) algorithm (*56*) (*SI Appendix*, Tables S1 and S2). We identified single putative Chlamydomonas orthologs for most Arabidopsis TOC components, as well as some TIC components. Because genes with similar functions are more likely to be co-expressed than random genes (*57–59*), we calculated their associated Pearson’s correlation coefficients (PCC) as a measure of transcriptional correlation (Fig. 1). We used the 26S proteasome as a control for a strong positive correlation (Fig. 1 A and C) (*59*). A pairwise comparison of all Chlamydomonas genes provided a control for lack of correlation (Fig. 1*C*, *SI Appendix*, Table S3). The analysis was based on RNAseq data from 518 samples derived from 58 independent experiments (for more details, see Materials and Methods). A positive PCC value indicates that two genes are co-expressed, a PCC value close to zero indicates no correlation, while a negative PCC value indicates that two genes have opposite expression patterns. We arranged genes in groups by hierarchical clustering to display their co-expression relationships (Fig. 1 *A* and *B*) (*SI Appendix*, Table S3). By comparison to the proteasome, the overall correlation of translocon components considered in this analysis is relatively weak (*SI Appendix*, Table S4).

**Fig. 1.**
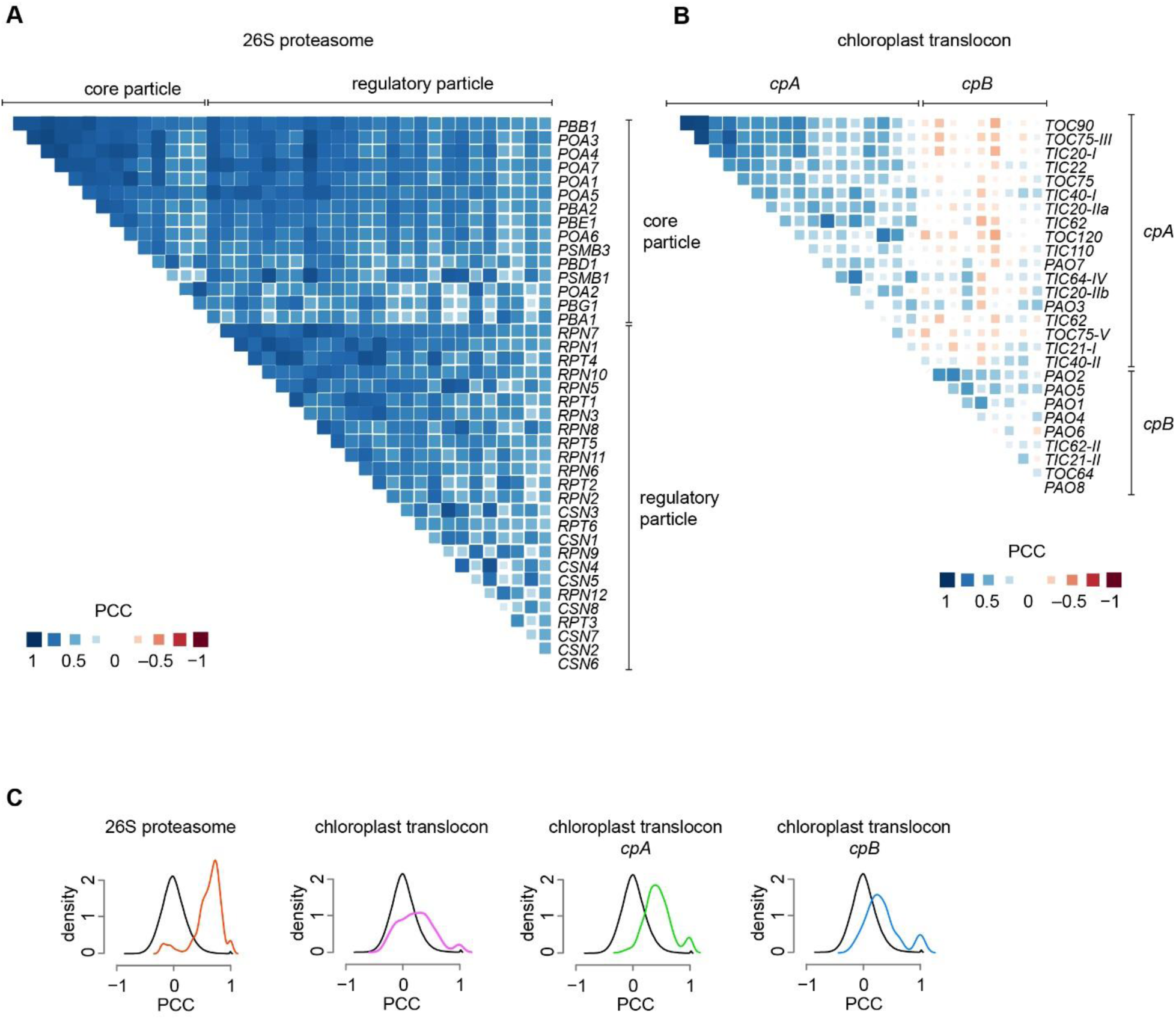
Co-expression patterns of Chlamydomonas genes encoding components of the plastid translocon. (*A*) Correlation matrix for Chlamydomonas genes encoding subunits of the 26S proteasome (listed in *SI Appendix*, Table S3). (*B*) Correlation matrix for Chlamydomonas genes encoding components of the chloroplast translocon identified through BLAST analysis (listed in *SI Appendix*, Table S3). (*C*) Distribution of Pearson’s correlation coefficients (PCCs) for all gene pairs encoding core and regulatory particles of the 26S proteasome (red line), and gene pairs for the chloroplast translocon components, together (magenta line) or as a function of their subgroup, *cpA* (green line) and *cpB* (blue line). PCC distribution for all gene pairs in the genome (black line) is shown to indicate the absence of correlation (i.e. negative control). Statistical significance of all distributions was tested by Kolmogorov-Smirnov test, comparing each sets of PCC values with a randomly generated normal distribution of equal element number. The results of this analysis are available in *SI Appendix*, Table S4.

Nevertheless, we found that Chlamydomonas chloroplast translocon genes can be classified in two clusters, hereafter referred to as *cpA* and *cpB* (Fig. 1 *B* and *C*) (*SI Appendix*, Table S3). The most strongly correlated genes in c*pA* include *TOC120*, *TOC90*, *TOC75*, *TIC110*, *TIC20*, previously shown to be essential for protein import in Arabidopsis (*11, 29, 30*). Except for *TOC64*, the much less correlated genes in *cpB* encode paralogs of genes in *cpA*. We surmise that the subgroup distribution may indicate a functional separation of components that are essential core translocon components (*cpA*) from those that may be dispensable or specialized (*cpB*). We obtained very similar results when performing an identical analysis with Arabidopsis plastid translocon genes (*SI Appendix,* Fig. S1), recapitulating previously reported expression patterns (*60*).

The BLAST searches identified all known subunits of the Chlamydomonas TOC complex (*SI Appendix*, Table S2). By contrast, similar BLAST searches failed to uncover all Chlamydomonas orthologs for the recently characterized components of the Arabidopsis TIC complex, except for *TIC20,* yet Chlamydomonas orthologs for Tic56 and Tic214 were previously proposed (*44*). Thus, to determine the composition of the Chlamydomonas TIC complex, we tagged Chlamydomonas Tic20 with a C-terminal yellow fluorescent protein (YFP) and a triple FLAG epitope (*61*) and carried out an immunopurification assay under native conditions, followed by mass-spectrometric analysis.

This strategy identified three Tic20-interacting proteins (Table 1, *SI Appendix*, Table S6), whose structural domain organization is similar to that of Arabidopsis Tic56, Tic100, and Tic214, respectively (*SI Appendix*, Fig. S2). Following the Arabidopsis naming convention, we will refer to these proteins as Chlamydomonas Tic56, Tic100, and Tic214. Tic56 and Tic100 are encoded by the nuclear genes Cre17.g727100 and Cre06.g300550, respectively, while Tic 214 is encoded by the essential chloroplast gene *orf1995* (*62*), hereafter referred to as *tic214*.

**Table 1.**
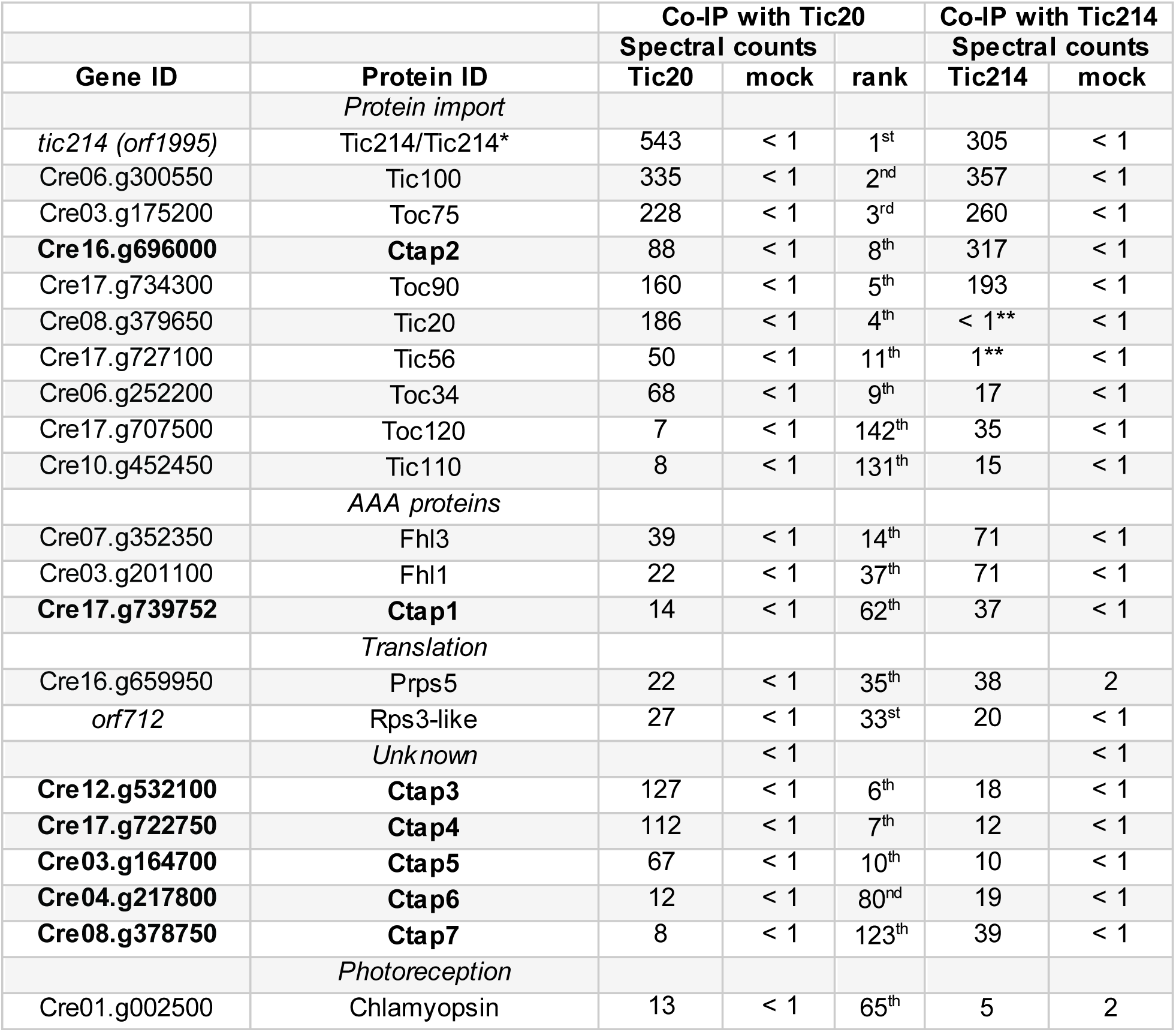
Spectral counts for proteins co-immunoprecipitated with Tic20 and Tic214. Tic214* denotes the 110 kDa protein band detected by the Tic214 antibody. The newly identified <u>c</u>hloroplast translocon <u>a</u>ssociated <u>p</u>roteins (Ctap) are highlighted in bold. Tic214 interacting partners were eluated under denaturing conditions and gel regions corresponding to the antibody chains were excluded during HPLC-MS/MS analysis. The asterisks (**) indicate those proteins that escaped detection because of their co-migration with the antibody chains.

Remarkably, several other proteins also co-immunoprecipitated with Tic20, including orthologs of the Arabidopsis TOC: Toc90 (Cre17.g734300), Toc120 (Cre17.g707500), Toc75 (Cre03.g175200) and Toc34 (Cre06.g252200) (Table 1). These findings suggest that the Chlamydomonas TIC and TOC complexes form a stable supercomplex (here referred to as TicToc) that spans the outer and inner chloroplast membranes, as previously shown in land plants (*9, 25, 63*).

We next performed a reciprocal co-immunoprecipitation followed by mass-spectrometry, using Tic214 as bait to confirm that the TicToc supercomplex indeed comprises chloroplast-encoded Tic214. Since the chloroplast genome can be manipulated with relative ease by homologous recombination in Chlamydomonas (*64*), we inserted three copies of the HA tag into the endogenous *tic214* locus (*SI Appendix*, Fig. S3*A*). Immunoblot analysis using an anti-HA antibody or an antibody raised against Tic214 detected a protein of smaller size than expected (around 110 kDa, hereafter referred to as Tic214*) (*SI Appendix*, Fig. S3*B*). Nevertheless, we retrieved peptides covering most of the Tic214 protein sequence upon affinity purification (*SI Appendix*, Fig. S3 *C* and *D*, Table S7). This result confirms that Tic214 is translated over the entire gene length and suggests that its abnormal electrophoretic mobility arises from at least one proteolytic nick, which may be introduced in the cell or during sample preparation, resulting in two or more stable fragments that remain part of the complex.

As expected from its role as bait in the immunoprecipitation, Tic214 emerged as the protein with the highest number of spectral counts (Table 1, *SI Appendix*, Table S7). Importantly, the analysis confirmed that Tic214 interacts with the same TOC subunits found in association with Tic20: namely Toc34, Toc75, and Toc90 (Table 1). We did not detect peptides derived from Tic20 or Tic56, likely because their electrophoretic mobility overlap with those of the antibody chains migrating in gel regions that were excluded from the analysis.

Taken together, these results suggest that the molecular architecture of the TOC and TIC complexes of land plants are conserved in chlorophyte algae, despite the lack of strong enough sequence similarity of some of its components that would have allowed their detection by BLAST analysis. Moreover, the ability of TIC and TOC to form a TicToc supercomplex, as well as the presence of a single chloroplast-encoded protein (Tic214), are phylogenetically conserved.

### Several novel proteins associate with the TicToc supercomplex

Aside from the known TOC and TIC subunits discussed above, we identified several uncharacterized proteins in our pull-downs using Tic20 (*SI Appendix*, Table S6) and Tic214 (*SI Appendix*, Table S7) as bait. The overlapping set of proteins found in both pull-downs contained three FtsH-like AAA proteins (Fhl1, Fhl3, and an uncharacterized <u>Ft</u>sH-like protein encoded by Cre17.g739752, that we named Ctap1 for <u>c</u>hloroplast translocon <u>a</u>ssociated <u>p</u>rotein 1). These three proteins lack the zinc-binding motif that is critical for the catalytic activity of typical FtsH metalloproteases; in this respect, they strongly resemble the Arabidopsis FtsHi proteins that were recently shown to associate with translocating pre-proteins during *in vitro* import reactions and serve as ATP-driven import motors (*50*). The immunoprecipitations also identified six additional previously uncharacterized proteins (Ctap2-7; encoded by Cre16.g696000, Cre12.g532100, Cre17.g722750, Cre03.g164700, Cre04.g217800 and Cre08.g378750, respectively) (Table 1, *SI Appendix*, Tables S6 and S7) that co-purified with both Tic20 and Tic214. Ctap2 is a 150 kDa protein that contains a C-terminal domain with homology to UDP-N-acetylglucosamine-pyrophosphorylases, while Ctap3-7 have no recognizable sequence motifs.

Fhl1, Ctap1, and Ctap5 bear predicted chloroplast transit peptides according to the chloroplast localization prediction software Predalgo (*65*), and Fhl3 and Ctap6 contain predicted transmembrane regions that may anchor them in the chloroplast envelope. A recently published genome-wide algal mutant library (*66*) does not contain any insertional mutants that would disrupt the coding region of any of these uncharacterized genes, hinting that these genes may be essential.

### The expression of TicToc components is coordinated across compartmental boundaries

As an extension of the approach described in Fig. 1, we surmised that if the newly identified proteins in the Tic20 and Tic214 pull-downs represented genuine components of the TicToc supercomplex, their encoding genes likewise might be co-expressed. To test this hypothesis, we recalculated the PCC correlation matrix after the addition of these genes to our original list of chloroplast translocon components (Fig. 2). Indeed, all genes coding for the proteins that co-purified with Tic20 and Tic214 exhibited correlated expression, as shown in Fig. 2 *A* and *C* (group *cp2*) (*SI Appendix*, Table S4).

**Fig. 2.**
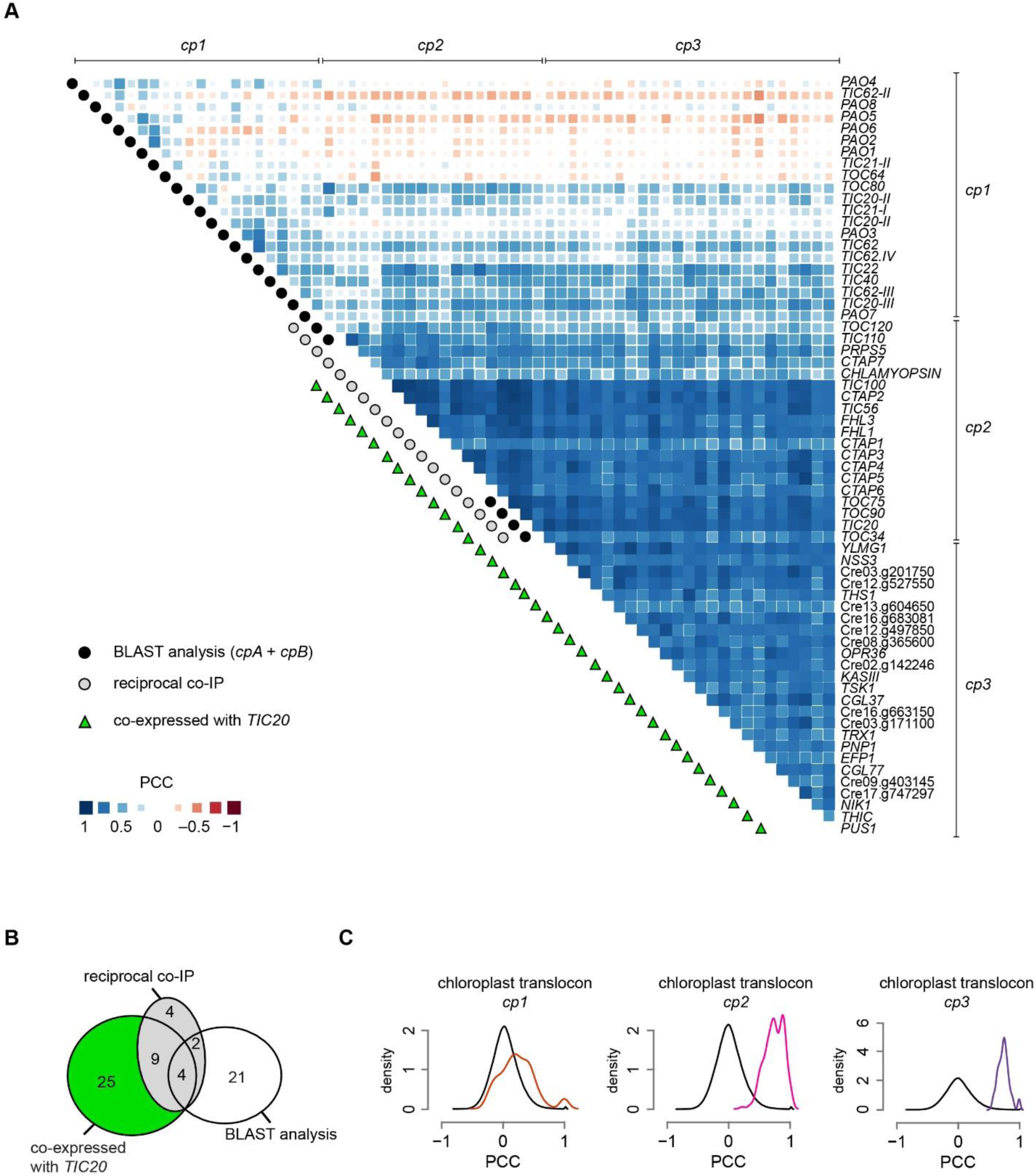
Co-expression patterns of known and newly identified chloroplast translocon components in Chlamydomonas. (*A*) Correlation matrix of chloroplast translocon genes from subgroups *cp1* (comprising most of *cpA* and *cpB* genes shown in Fig. 1A), subgroup *cp2* (containing genes identified during both Tic20 and Tic214 pull-downs and listed in Table 1) and subgroup *cp3* (comprising genes only co-expressed with *TIC20* and listed in *SI Appendix*, Table S8). (*B*) Venn diagram related to gene subgroups shown in panel A. (*C*) PCC distributions for genes belonging to subgroups *cp1* (red line), *cp2* (magenta line), and *cp3* (purple line) and for all gene pairs in the genome (black line) used as negative control. Statistical significance of all distributions was tested by Kolmogorov-Smirnov test, comparing each sets of PCC values with a randomly generated normal distribution of equal element number. The results of this analysis are available in *SI Appendix*, Table S4.

Based on these results, we next extended the co-expression analysis to look for other genes that correlate in their expression with *TIC20*. In this way, we identified 25 additional genes exhibiting co-expression with *TIC20* (*SI Appendix*, Table S8), (group *cp3,* Fig. 2 *A-C*). Since proteins encoded by *cp3* genes were not identified in the pull-downs, we hypothesize that they may be involved in the regulation/assembly of the translocon, or be more loosely associated and washed off during affinity purification. We did not further pursue these proteins in this investigation.

Because the vast majority of the expression datasets used in our analyses are based on polyA-selected RNA samples, they do not include mRNAs transcribed inside an organelle where polyA addition does not occur. Thus, to test for co-expression between *tic214* and nucleus-encoded TIC and TOC genes, we used a dataset that was generated by random priming and interrogated nuclear and chloroplast gene expression changes over the diurnal cycle (12 h dark / 12 h light regime) (*67*). We observed that the expression profile of *tic214* followed that of the nucleus-encoded TIC and TOC components closely, with peaks in the early dark and light phases (Fig. 3*A*). The expression of *tic214* also correlated with that of *orf2971*, a yet uncharacterized chloroplast gene. *orf2971* encodes a protein that resembles Ycf2, the chloroplast-encoded subunit of the Ycf2/FtsHi protein import motor recently identified in Arabidopsis (*50*). Orf2971 was identified during the Tic20 pull-down (*SI Appendix*, Table S6).

**Fig. 3.**
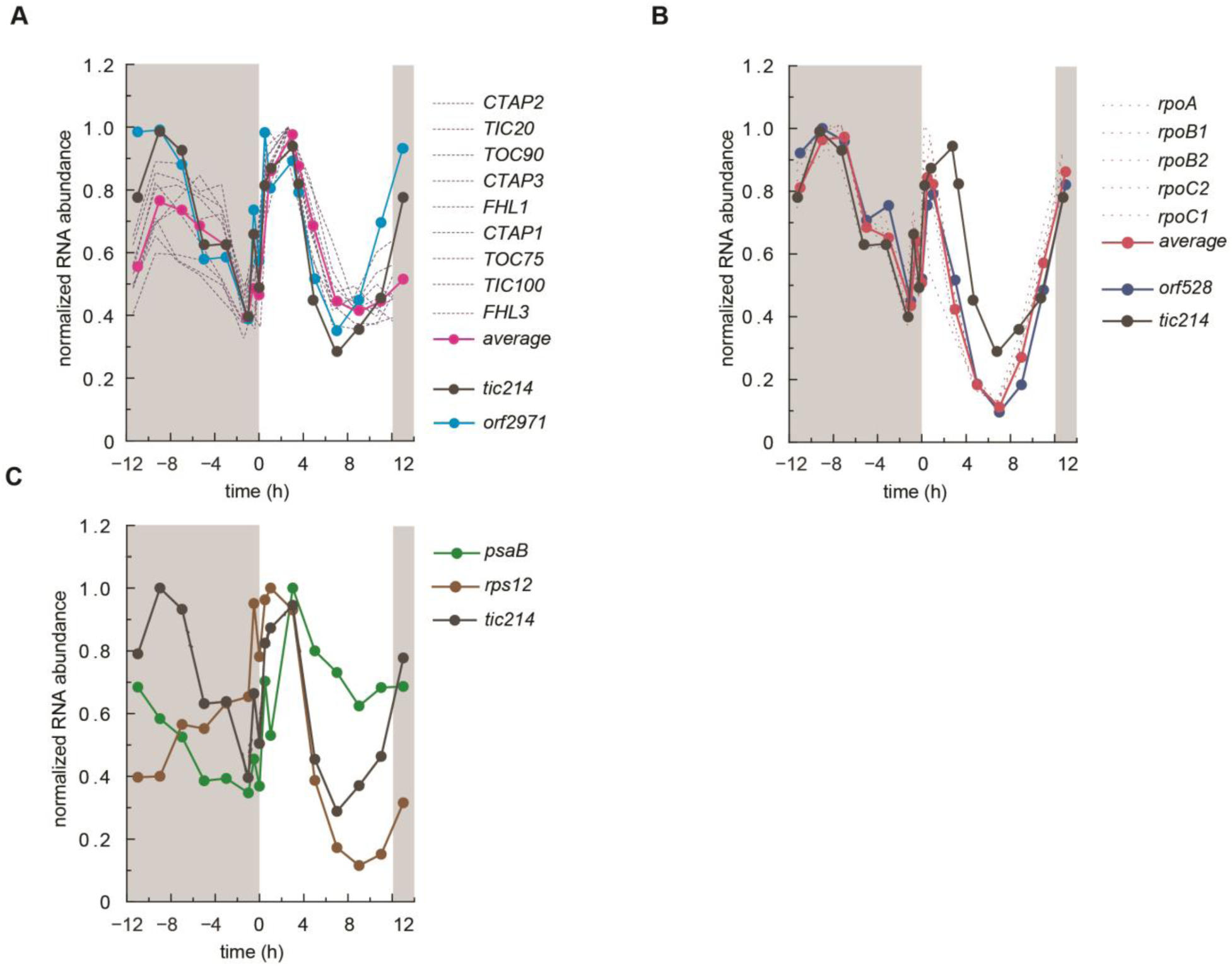
Co-expression profiles of chloroplast-encoded and nucleus-encoded translocon components over the course of a diurnal cycle in Chlamydomonas. (*A*) *tic214* (*orf1995*) (black) and *orf2971* (light blue), and co-expressed nucleus-encoded translocon components (blue). (*B*) *tic214* (black), *orf528* (blue) and co-expressed plastid-encoded RNA polymerase genes (PEP) (dark red). (*C*) *tic214* (black), *psaB* (green) and *rps12* (brown).

A similar expression pattern was also observed for the chloroplast *rpo* genes (encoding the chloroplast RNA polymerase subunits) and *orf528*, another chloroplast gene of unknown function (*68*) (Fig. 3*B*). Although unexpected, these correlations are specific, as we observed no correlation between the expression profiles of *tic214* and that of most other chloroplast genes, including those involved in photosynthesis and ribosome biogenesis, such as *psaB* and *rps12,* which thus serve as negative controls (Fig. 3*C*).

Taken together, the results of the Tic20 and Tic214 co-immunoprecipitation, combined with co-expression analysis, paint a comprehensive picture of translocon composition that strongly supports the assignment of the newly identified proteins as *bona fide* subunits and/or potential biogenesis factors and regulators of the TicToc supercomplex Chlamydomonas. Moreover, our analysis points to an extensive control network(s) that coordinates their expression across compartmental boundaries and over the diurnal cycle.

### The TicToc supercomplex is stable

To assess the stability of the TicToc supercomplex, we affinity-purified the Tic20-tagged complex and analyzed it by blue native polyacrylamide gel electrophoresis (PAGE), followed by SDS-(2D blue native/SDS-PAGE). We found that the proteins associated with Tic20 co-migrate as a large complex (Fig. 4 *A* and *B*). These spots were not detected by silver staining when the affinity purification was performed using extracts derived from an untagged strain (*SI Appendix*, Fig. S4*A*). Mass-spectrometry of discrete protein spots derived from the large complex identified Tic56 and Tic214, as well as Toc34, Toc75, Toc90, Ctap2, Ctap4 and Ctap5. Moreover, we obtained independent confirmation about the presence of Tic214, Tic20, Tic56 and Tic100 by targeted immunoblot analysis with antibodies raised against each protein (Fig. 4*B*, *SI Appendix*, Fig. S4*B*).

**Fig. 4.**
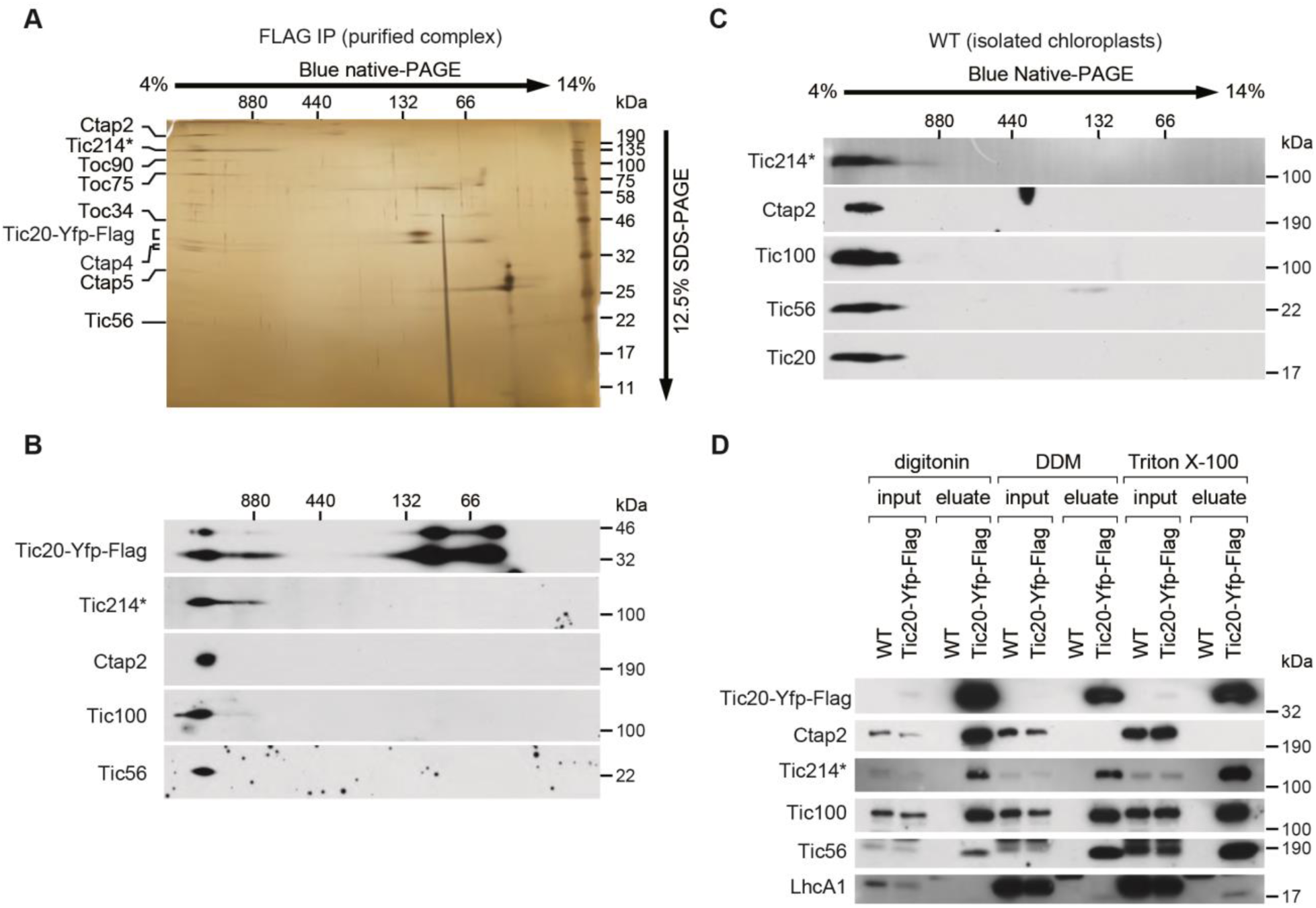
Tic20 is part of a stable chloroplast translocon supercomplex in Chlamydomonas. (*A*) 2D blue native/SDS-PAGE separation of the purified Tic20-YFP-FLAG_3x_ fraction (in the presence of digitonin) followed by silver staining. Proteins identified by mass-spec analysis are indicated. (*B*) Proteins were separated as in panel A and analyzed by immunoblotting with the indicated antibodies. (*C*) Isolated chloroplasts were solubilized with digitonin and analyzed as in panel A. (*D*) Purification of the Tic20-YFP-FLAG_3x_ containing supercomplex was carried out after solubilization with digitonin, dodecylmaltoside (DDM), or Triton X-100. The purified fractions (eluate) together with 1% of the input fractions used for the purification (input) were analyzed by SDS-PAGE and immunoblotting with the indicated antibodies. Mock-purified fractions (WT) were also analyzed. Tic214* in panels *A*, *B*, *C,* and *D* denotes the 110 kDa protein band detected by the Tic214 antibody.

As a control, we solubilized the TicToc supercomplex from chloroplast membranes derived from a strain lacking the Tic20-tagged protein. Immunoblot analysis after 2D blue native/SDS-PAGE revealed that Tic20, Tic56, Tic100, and Tic214 all migrated as part of the large complex (Fig. 4*C*), demonstrating that the presence of a FLAG-tag has no impact on the formation or composition of the TicToc supercomplex.

Next, we repeated the Tic20-FLAG affinity purification from membrane extracts prepared with different non-ionic detergents: digitonin, dodecylmaltoside (DDM), and Triton X-100. Immunoblotting analysis after 2D blue native/SDS-PAGE (Fig. 4*D*) detected the four TIC complex components (Tic20, Tic56, Tic100, and Tic214), following membrane solubilization with all detergents, confirming these proteins as core components of the chloroplast translocon. By contrast, Ctap2 was only found with Tic20 in the presence of digitonin, but not with DDM or Triton X-100 (Fig. 4*D*), suggesting that its association is detergent-sensitive.

### The TicToc supercomplex functionally associates with chloroplast pre-proteins

To validate that the TicToc supercomplex functions in importing chloroplast-targeted proteins, we produced two small pre-proteins in *E. coli*: pre-Rubisco small subunit Rbcs2 and pre-ferredoxin Fdx1, bearing purification tags and TEV protease cleavage sites (Fig. 5*A*). We then incubated the purified pre-proteins with intact Chlamydomonas chloroplasts in the presence of ATP (0.3 or 3 mM) to initiate the import reaction. After a 15 min incubation, we collected chloroplasts and any bound pre-proteins (“Tot” in Fig. 5*B*) by centrifugation. An aliquot of the enriched chloroplast fraction was subjected to freeze-thaw cycles to extract fully imported proteins from the stroma (“Sup” in Fig. 5*B*). For both pre-proteins, we detected the precursor and the mature cleaved protein in the Tot fraction, whereas the stromal fraction (Sup) was enriched with mature cleaved proteins. In the case of pre-Fdx1, all import events (binding of precursor, translocation, and maturation) were dependent on externally added ATP (Fig. 5*B*); for pre-Rbcs2, we observed mature size proteins in the Sup fraction even in the absence of added ATP, possibly mediated by residual ATP contained in the isolated chloroplasts (Fig. 5*B*).

**Fig. 5.**
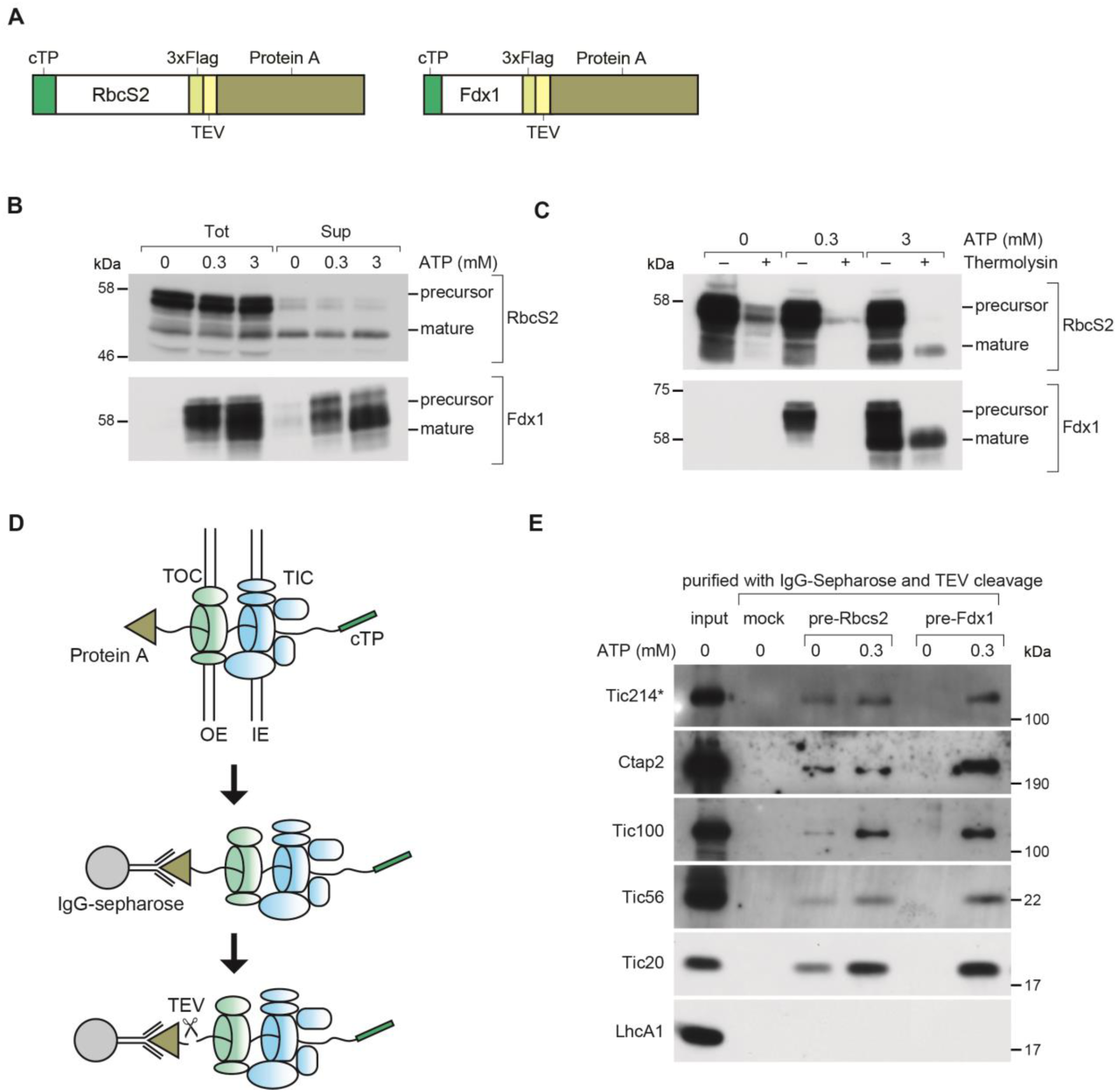
*In vitro* import assay of chloroplast pre-proteins and purification of translocation intermediates in Chlamydomonas. (*A*) Schematic representation of the two chloroplast pre-proteins used for *in vitro* import assays and the purification of translocation intermediates. (*B*) Chloroplast pre-proteins were incubated with intact chloroplasts isolated from Chlamydomonas in the presence of the indicated concentrations of ATP. Chloroplasts were re-isolated, washed, and directly analyzed (“Tot”) or fractionated into soluble fraction containing stroma (“Sup”). (*C*) Same assay as in panel B, except that chloroplasts were treated with thermolysin after protein import. In (*B)*, a band corresponding to the mature form of RbcS2 is detected in the absence of ATP and recovered in the Sup fraction. This band does not represent fully translocated protein but is most likely formed or released during the freeze-thaw cycles used for chloroplast lysis. This notion was verified in (*C*) with thermolysin treatment, which shows that there is no fully translocated mature RbcS2 in the absence of ATP. The RbcS2 import experiment was repeated several times with similar band profiles. (*D*) Outline of the method used to isolate chloroplast intermediate translocation complexes. After import, washed chloroplasts were solubilized with digitonin, and translocation intermediates were purified with IgG-Sepharose and eluted by TEV protease cleavage. Mock purification was carried out without the addition of pre-proteins. IE, OE, inner, outer envelope membrane, respectively. (*E*) Purified fractions shown in panel D were analyzed by SDS-PAGE and immunoblotting with the indicated antibodies. Tic214* denotes the 110 kDa protein band detected by the Tic214 antibody.

To confirm the ATP dependence of pre-protein translocation into the stroma, we used the protease thermolysin (*69*), which degrades proteins associated with the outer chloroplast membrane, but cannot access those translocated into the stroma. In this assay, the addition of ATP was required for complete translocation and maturation for both pre-Rbcs2 and pre-Fdx1 (Fig. 5*C*, *SI Appendix*, Fig. S5).

To identify the TIC and TOC components that are juxtaposed to the pre-proteins during import, we isolated chloroplasts at the end of an import reaction carried out with or without ATP addition. We then solubilized membrane fractions with digitonin and purified translocation intermediates through the protein A tag by affinity chromatography on IgG Sepharose followed by TEV-mediated elution under non-denaturing condition as depicted in Fig. 5*D*. As a negative control, we omitted the pre-incubation step with protein A-tagged precursors. When probing the eluted fractions with antibodies against TIC complex components Tic20, Tic56, Tic100, and Tic214, we found that the association of all TIC proteins was stimulated by ATP (Fig. 5*E*), consistent with an energized translocation event even for pre-RbcS2. Ctap2 was among the proteins identified in the eluted fractions (Fig. 5*E*), providing further evidence of its potential role in pre-protein import.

We confirmed these results by unbiased mass spectrometry of the immunoprecipitated complexes. In the case of the Fdx1 precursor translocation intermediate, several TIC and TOC proteins were enriched in the purified fraction in an ATP-dependent manner. Notably, Tic214 and Ctap2 topped the list with the highest spectral counts detected in the presence of ATP (*SI Appendix*, Fig. S6 *A* and *B*), followed by other proteins identified above (Table 2). The same proteins also associated with pre-Rbcs2. However, in this case, the spectral counts were comparable in the presence or absence of ATP (*SI Appendix*, Table S9). This result is consistent with the *in vitro* import assays shown in Fig. 5 *B* and *E*.

**Table 2.**
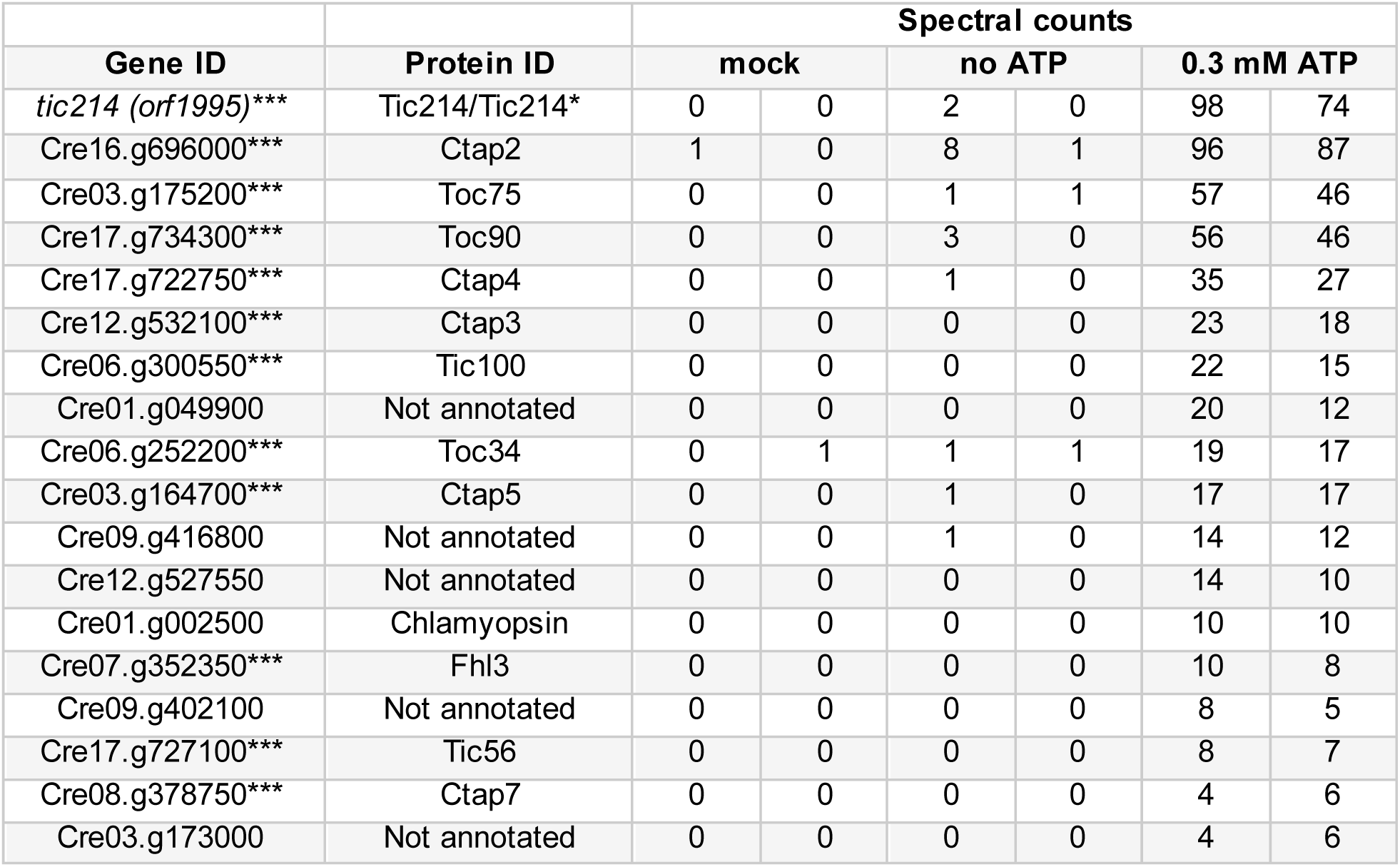
Spectral counts for proteins associated with pre-Fdx1 translocation intermediates. The three asterisks (***) indicate genes co-expressed with *TIC20*. Tic214* denotes the 110 kDa protein band detected by the Tic214 antibody.

| Gene ID | Protein ID | Spectral counts |  |  |  |  |  |
| --- | --- | --- | --- | --- | --- | --- | --- |
|  |  | mock |  | no ATP |  | 0.3 mM ATP |  |
| *tic214 (orf1995)* *** | Tic214/Tic214* | 0 | 0 | 2 | 0 | 98 | 74 |
| Cre16.g696000*** | Ctap2 | 1 | 0 | 8 | 1 | 96 | 87 |
| Cre03.g175200*** | Toc75 | 0 | 0 | 1 | 1 | 57 | 46 |
| Cre17.g734300*** | Toc90 | 0 | 0 | 3 | 0 | 56 | 46 |
| Cre17.g722750*** | Ctap4 | 0 | 0 | 1 | 0 | 35 | 27 |
| Cre12.g532100*** | Ctap3 | 0 | 0 | 0 | 0 | 23 | 18 |
| Cre06.g300550*** | Tic100 | 0 | 0 | 0 | 0 | 22 | 15 |
| Cre01.g049900 | Not annotated | 0 | 0 | 0 | 0 | 20 | 12 |
| Cre06.g252200*** | Toc34 | 0 | 1 | 1 | 1 | 19 | 17 |
| Cre03.g164700*** | Ctap5 | 0 | 0 | 1 | 0 | 17 | 17 |
| Cre09.g416800 | Not annotated | 0 | 0 | 1 | 0 | 14 | 12 |
| Cre12.g527550 | Not annotated | 0 | 0 | 0 | 0 | 14 | 10 |
| Cre01.g002500 | Chlamyopsin | 0 | 0 | 0 | 0 | 10 | 10 |
| Cre07.g352350*** | Fhl3 | 0 | 0 | 0 | 0 | 10 | 8 |
| Cre09.g402100 | Not annotated | 0 | 0 | 0 | 0 | 8 | 5 |
| Cre17.g727100*** | Tic56 | 0 | 0 | 0 | 0 | 8 | 7 |
| Cre08.g378750*** | Ctap7 | 0 | 0 | 0 | 0 | 4 | 6 |
| Cre03.g173000 | Not annotated | 0 | 0 | 0 | 0 | 4 | 6 |

Taken together, our results show that the components of the TicToc supercomplex, identified by bioinformatics, protein pull-downs, co-expression analysis, are indeed part of an import-competent machinery.

### *tic214* expression is required for normal chloroplast morphology and proteostasis

To assess the role of *tic214 in vivo*, we engineered a strain (Y14) that allows conditional repression of this chloroplast gene in the presence of vitamins (vitamin B_12_ and thiamine, hereafter referred to as “Vit”) (Fig. 6*A*, *SI Appendix*, Fig. S7) (*70–72*). As a control, we used the parental strain (A31), in which the addition of Vit does not affect *tic214* expression. As expected for an essential gene, Vit addition to the medium blocked the growth of Y14 cells but not of A31 control cells (Fig. 6 *B* and *C*). Immunoblot analysis confirmed a time-dependent decrease in the level of Tic214* (Fig. 6*D*). This result further validated that Tic214* originates from *tic214*.

**Fig. 6.**
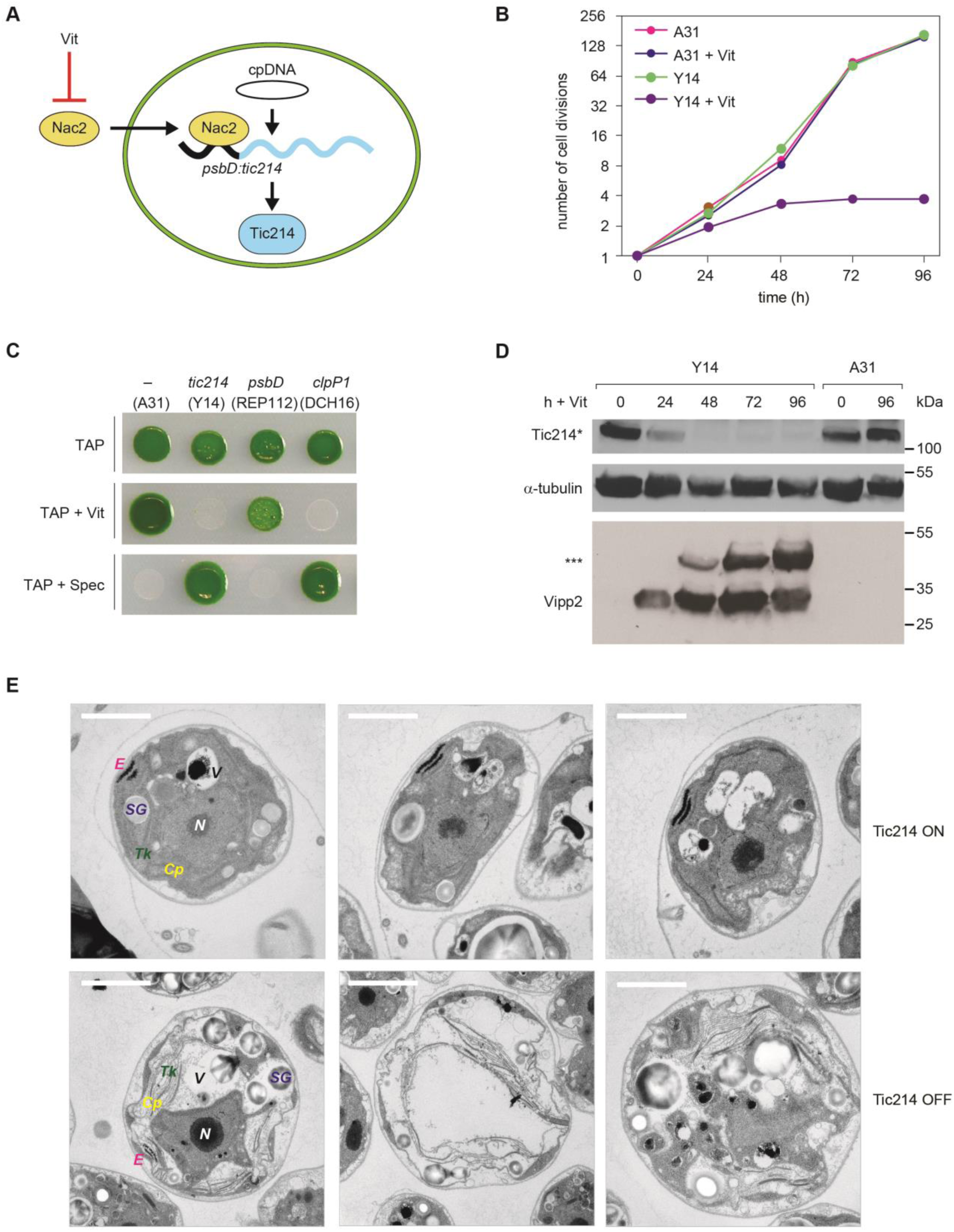
Conditional depletion of Tic214 inhibits cell growth. (*A*) Schematic illustration of selective Vit-mediated Tic214 depletion in Y14 cells. The Nac2 protein is translated in the cytosol and imported into the chloroplast, where it is required for expression of the chimeric *psbD:tic214* transgene. Accumulation of the Nac2 protein, and in turn of Tic214, is down-regulated upon the addition of thiamine and vitamin B_12_ to the growth media (“Vit”). (*B*) Growth curve of Y14 and the A31 control strains in TAP without or with Vit (20 μg /L vitamin B_12_ and 200 μM thiamine HCl) under standard light conditions (∼60 μmol m^-2^s^-1^ irradiance). (*C*) Growth of A31, Y14, REP112 and DCH16 strains was tested by spotting cells on plates containing acetate (TAP) with and without Vit (20 μg/L vitamin B_12_ and 200 μM thiamine HCl) and in TAP supplied with 100 μg/ml spectinomycin (Spec) on which only chloroplast transformants can survive. The chloroplast gene controlled by Nac2 in each strain is indicated at the top. A31 is used as a control, since no chloroplast transcript is under the control of the Vit-mediated Nac2 expression in this strain. Expression of the chimeric *psbD:tic214* gene is repressed by Vit in Y14, expression of the endogenous *psbD* is repressed by Vit in REP112 whereas the chimeric *psbD:clpP1* is repressed by Vit in DCH16 (*70, 71*). (*D*) Immunoblot analysis of Tic214 and Vipp2 in Y14 and A31 treated with Vit for the indicated times. The three asterisk indicate a potential SDS-resistant Vipp2 aggregate. Tic214* denotes the 110 kDa protein band detected by the Tic214 antibody. α-tubulin was used as a loading control. (*E*) Electron microscopy of Y14 cells supplemented with Vit for 96 hours (lower row, “Tic214 OFF”). The control cells without Vit treatment (“Tic214 ON”) are shown in the upper row (scale bar = 2 µm). Intracellular compartments are labeled as follows: *Cp* = chloroplast; *Tk* = thylakoids; *E* = eyespot; *SG* = starch granules; *V* = vacuole; *N* = nucleus.

To further characterize the consequences of Tic214 depletion, we visualized cells by transmission electron microscopy. As shown in Fig. 6*E*, *tic214* repression caused a massive cellular swelling and resulted in a chloroplast with highly disorganized thylakoid ultrastructure and accumulation of starch granules, both typical traits of cells experiencing chloroplast proteotoxic stress (*71, 73, 74*). In agreement with this observation, we found that the time-dependent decrease in Tic214 inversely correlates with the induction of Vipp2 (Fig. 6*D*), a marker of the chloroplast unfolded protein response (cpUPR) (*71, 75*). The cpUPR is a stress-induced signaling network that responds to impairment of chloroplast proteostasis to maintain organellar health (*71, 73, 76-79*).

### *tic214* repression impairs protein import into chloroplasts

If Tic214 is a *bona fide* and essential component of the chloroplast translocon, its depletion should block protein import. To obtain direct evidence for translocation defects, we aimed to detect an accumulation of non-translocated chloroplast pre-proteins. To this end, we used tandem mass spectrometric analysis of protein extracts from strain Y14 collected following Tic214 repression and compared the results to the A31 control strain (*SI Appendix*, Table S10). Since unimported chloroplast proteins are quickly degraded by the cytosolic ubiquitin-26S proteasome system (*16, 80, 81*), we incubated each culture with the proteasome inhibitor MG132 before sampling. We limited our analysis to those proteins for which at least ten peptides could be identified in one of the six conditions used in the experiment (2, 4, and 6 days; -/+Vit). This arbitrary cut-off narrowed the dataset to 1,458 of the 5,105 proteins detected, of which 1,427 are encoded by nuclear genes, and 427 are predicted to be chloroplast-localized (*SI Appendix*, Table S11). Among these, we identified 44 proteins for which we detected sequences derived from their predicted transit peptides only upon Tic214 repression (Fig. 7 *A* and *B*, *SI Appendix*, Table S11), indicating that their translocation into the chloroplast was impaired. Some of the peptides included the transit peptide cleavage site, suggesting that they were derived from pre-proteins that did not access the stromal presequence protease (*82, 83*). In addition, seven of these proteins were acetylated at their N-terminal methionine, and five were ubiquitylated only upon Tic214 repression (*SI Appendix*, Table S11). Together, the presence of peptides containing chloroplast transit peptide sequences, the accumulation of uncleaved pre-proteins, and the detection of two post-translational modifications observed only in cytosolic proteins (*36, 84*), strongly suggests that the import of these chloroplast proteins is impaired in the absence of Tic214.

**Fig. 7.**
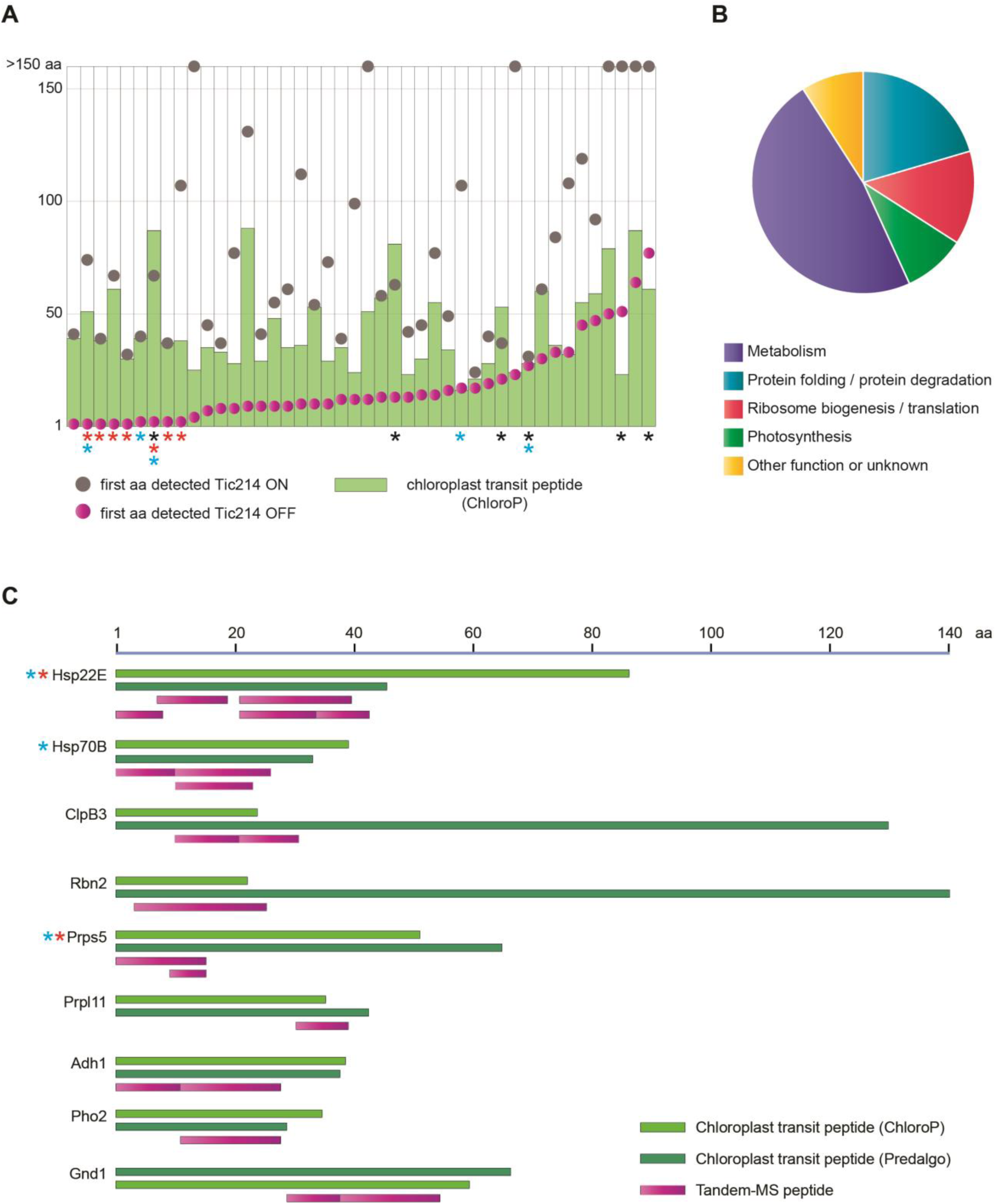
Detection of chloroplast precursor proteins upon Tic214 depletion. (*A*) Schematic representation of the 44 proteins (listed in *SI Appendix*, Table S11) for which one or more peptide covering the putative chloroplast transit peptide (cTP) could be detected, selectively upon Tic214 depletion. The green bar indicates the length of the cTP (i.e. number of amino acids) as predicted by ChloroP (*96*). The circles indicate the first amino acid position of the most N-terminal MS-peptide detected in the presence (grey) or absence (magenta) of Tic214. The black asterisks indicate those cases when peptides containing precursor sequences were observed only according to the cTP length predicted by Predalgo (*65*). The red and light blue asterisks indicate proteins that have an acetylation on their N-terminal methionine and are ubiquitylated upon Tic214 repression, respectively. (*B*) Pie-chart summarizing the functional classification of the proteins shown in A. Metabolism (n=21); Protein folding / Protein degradation (n=9); Ribosome biogenesis / translation (n=6); Photosynthesis (n=4); Other functions or unknown (n=4). (*C*) The location and length of peptides spanning the cTP are depicted for some of the proteins shown in A. The light and dark green bars indicate the cTP length as predicted by ChloroP and Predalgo, respectively. The magenta bars indicate the length of each MS-peptide, selectively detected upon Tic214 depletion.

A manually-curated functional annotation of these 44 proteins revealed that they are involved in various metabolic pathways, including some important reactions such as amino acid synthesis, purine synthesis, one-carbon metabolism, protein folding/degradation, ribosome biogenesis and translation, and photosynthesis (Fig. 7 *B* and *C* and *SI Appendix*, Tables S11 and S12). These data suggest that the import of some chloroplast proteins with essential functions in chloroplast protein folding and ribosome biogenesis requires Tic214, thus explaining why Tic214 depletion causes chloroplast proteotoxicity, inducing the cpUPR. A recent study identified a set of 875 chloroplast stress-responsive proteins (*75*). Although we only detected 91 of these proteins by mass spectrometry here, a vast majority (78 out of 91) was upregulated upon Tic214 depletion, and the expression of 45 of the encoding genes is dependent on Mars1, a critical component of cpUPR signaling (*SI Appendix*, Fig. S8 and Table S13) (*75*).

## Discussion

Despite the universal importance of chloroplasts in sustaining life on Earth, there is surprisingly little consensus regarding the molecular machineries that carry out protein import into these organelles. Here, we demonstrate that all of the subunits of the 1-MDa TIC complex previously identified in Arabidopsis are functionally conserved in Chlamydomonas, indicating that this protein import module has been maintained over approximately 1.1 billion years of evolution (*85*). This conclusion is supported by a combination of gene co-expression, protein-protein interaction, and translocation analyses, yet has previously been controversial because sequence comparison analyses failed to identify orthologs for some components of the complex. Thus, our data show that proteins performing essential functions such as chloroplast protein import can be highly divergent in their primary sequence.

In agreement with their functional conservation, we found that in Chlamydomonas, as in Arabidopsis, the TIC complex interacts with the TOC complex to form a TicToc supercomplex. This supercomplex includes several uncharacterized proteins. In particular, we identified three AAA proteins (Fhl3, Fhl1, and Ctap1) and a potential Ycf2 ortholog, Orf2971, that may act as ATP-driven import motors, as previously shown for Arabidopsis and tobacco (*50*). In addition, we found six other uncharacterized chloroplast translocon-associated proteins (Ctap2-7) in the Chlamydomonas supercomplex that were not identified in Arabidopsis. Their topology, function, and evolutionarily conservation remain to be determined. Analysis of several Chlamydomonas transcriptomic datasets revealed that the expression patterns of genes encoding these newly identified components are highly correlated with known TIC and TOC subunits, supporting the idea that these proteins participate in the same biological process.

Our co-expression analysis uncovered additional nuclear and chloroplast genes encoding proteins that were not identified in co-immunoprecipitation experiments, yet may play a role in chloroplast translocon assembly or regulation. Elucidating the exact function of these genes will provide valuable insights into our understanding of the chloroplast protein import machinery.

Importantly, we found that the co-expression of translocon components is not restricted to nuclear-encoded subunits, for which co-regulation was previously shown in Arabidopsis (*60*), but also encompasses chloroplast-encoded subunits. Indeed, chloroplast *tic214* and *orf2971* are co-expressed with nuclear *TIC20* during the Chlamydomonas diurnal cycle, suggesting their common regulation. Thus, in addition to the photosynthetic complexes and the chloroplast ribosome (*86–88*), the chloroplast translocon emerges as yet another example of coordinated gene expression between the chloroplast and nuclear compartment.

In Arabidopsis, TIC and TOC are linked together through Tic236, an integral inner-membrane protein that directly binds to Toc75 via its C-terminal domain protruding into the intermembrane space (*89*). Although BLAST analyses identify an algal orthologue of Tic236 (Cre05.g243150), we did not detect this protein in our experiments or co-expression studies. Hence, how the TicToc supercomplex of Chlamydomonas is held together remains an open question.

To assess the role of the TicToc supercomplex *in vivo*, we engineered a conditional expression system for Tic214. Upon its depletion, we observed severe effects on chloroplast morphology and cell growth, as well as the induction of cpUPR target proteins. By shot-gun proteomic analysis *in vivo*, we positively identified tens of proteins that retained their chloroplast transit peptides upon Tic214 repression. These pre-proteins were detected in the presence of proteasome inhibitors, unmasking their otherwise transient nature. Since some of these proteins have essential functions, including those involved in chloroplast ribosome biogenesis, our data explain why Tic214 is indispensable for cell survival. Hence, the defect in chloroplast translation, previously observed in an Arabidopsis *tic56* mutant (*90*), is likely due to an indirect consequence of a malfunctioning TicToc supercomplex.

In conclusion, we here demonstrate the power of using Chlamydomonas as a model system to dissect the composition, evolution, and regulation of the chloroplast translocon, and, by extension, of other evolutionarily conserved multiprotein complexes. The outstanding stability of the TicToc supercomplex identified in this study, combined with the ease of growing large amounts of Chlamydomonas cells, now opens unprecedented opportunities for the structural and functional characterization of this essential molecular machinery that firmly rivets the inner and outer chloroplast envelope membranes.

## Materials and Methods

### Strains, Growth Conditions and Media

*Chlamydomonas reinhardtii* strains were grown on Tris-acetate phosphate (TAP) or minimal (HSM) solid medium containing 1.5% Bacto-agar (*91, 92*) at 25°C in constant light (60-40 μmol m^-2^ s^-1^), dim light (10 μmol m^-2^ s^-1^) or in the dark. Growth medium containing vitamin B_12_ and Thiamine-HCl (Vit) was prepared as previously described (*70*).

### Co-expression analysis

We mainly followed the steps outlined by (*93, 94*) to assess gene co-expression and generate *TIC20* co-expression networks. We collected raw reads from 58 independent RNAseq experiments, representing 518 samples, and re-mapped them to version v5.5 of the Chlamydomonas genome. Gene expression estimates, in fragments per kilobase of exon model per million reads mapped (FPKMs), were then normalized in three steps: 1) log_2_(FPKM + 1), to account for genes with zero FPKMs; 2) quantile normalization with the R package *preprocessCore*; 3) normalization by the mean of each gene across all samples, resulting in a scale-free dataset. For *A. thaliana*, we collected microarray data from AtGenExpress and processed them as described above. Gene lists were compiled from literature searches and BLAST results using the *A. thaliana* or *P. sativum* genes, and are given in SI Appendix, Tables S1 and S2. Co-expression was visualized with the R package *corrplot*, with the order of genes within each set (proteasome and chloroplast translocon) determined by hierarchical clustering using a combination of “hclust” and “FPC” methods in *corrplot*. The distribution of Pearson correlation coefficient (PCC) values for each gene set was plotted in R using the *density* function. To identify genes that are co-expressed with *C. reinhardtii TIC20*, we calculated the mutual ranks (MR) associated with all gene pairs (*93, 94*) and then converted MR values into network edge weights (an edge being the distance between two genes (or nodes)). For the highest stringency, we opted for the formula with the fastest rate of decay: edge = e^-(MR-1)/5^, and deemed a gene to be co-expressed with *TIC20* only if its network edge weight with *TIC20* was greater than 0.01.

### Construction of plasmids

#### Construction of pRAM73.19

A chloroplast integration plasmid carrying the *psbD* 5’UTR fused to the *tic214 (orf1995)* coding sequence was generated in two sub-cloning steps, as described below. First, 210 bp of the *psbD* promoter and 5’UTR and about 2.4 kbp of the *tic214* coding sequence were amplified from chloroplast genomic DNA using primer pair SR212/SR247 and SRSR246/SR231, respectively. These two PCR products were gel-purified, mixed in an equimolar amount, and used as a template for an overlap extension PCR by SR212/SR231. The resulting PCR product was gel-purified, digested by ClaI and EcoRI, and cloned in the chloroplast integration vector pUCatpXaadA digested with the same restriction enzymes. The resulting plasmid was verified by digestion analysis and named pRAM72. Next, about 2.7 kbp of the region upstream of the *tic214* 5’UTR was amplified from chloroplast genomic DNA using primer pair SR228/SR229. This PCR product was gel-purified, digested by SacI and XbaI, and cloned into pRAM72 digested with the same restriction enzymes. The resulting plasmid was verified by digestion analysis and DNA sequencing and named pRAM73.19. The sequence of the primers used during this cloning are reported below:

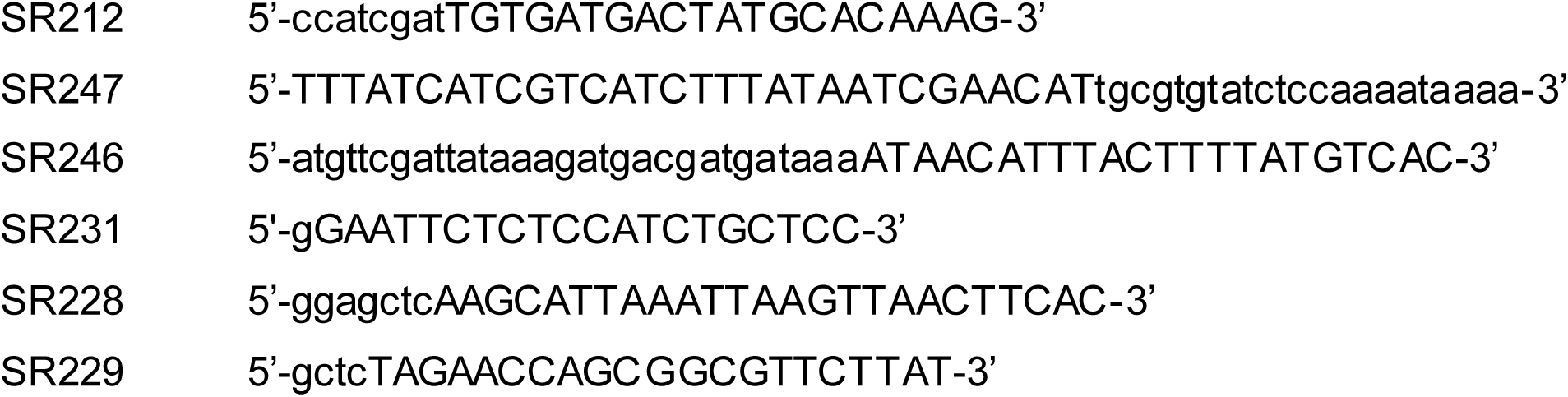

#### Construction of pED3 (to produce recombinant proteins used to raise the polyclonal antibody against Tic214)

A DNA fragment of *tic214* was amplified from genomic DNA using the primer pair ED5/ED6. This PCR product was gel-purified, digested by NcoI and XhoI, and cloned in the bacterial expression vector pET28a digested with the same restriction enzymes. The resulting plasmid was verified by digestion analysis and DNA sequencing and named pED3. The sequence of the primers used during this cloning are reported below:

ED5 5’-catgccatggTAAATGTAGCTAAACAAATATTA-3’

ED6 5’-ccgctcgagTGTTTTTCTCCAACGTAAAGT-3’

### Construction of the Y14 and NY6 strains

To generate the Y14 strain, the A31 strain (*70*) was transformed with the chloroplast integration plasmid pRAM73.19, where the *aadA* cassette (used as a selective marker) is located just upstream of the *psbD* 5’UTR fused to the *tic214* gene. To generate the NY6 strain, a WT strain was transformed with a chloroplast integration plasmid named Orf1995:HA (NdeI) (gift of E. Boudreau with the HA-11 epitope inserted in the 5’ coding sequence of the *tic214* gene. In this vector, *tic214* is under the control of its endogenous 5’ UTR, and the *aadA* cassette is adjacent to it. Chloroplast transformation was performed as described (*70*).

### Chloroplast isolation

The Chlamydomonas cell wall-deficient strains CC-400 (*cw-15* mt^+^) (a kind gift from H. Fukuzawa (Kyoto University), CC-4533 and CS1_FC1D12 (CC-4533 transformed with a *TIC20-YFP-FLAG_3x_* nuclear transgene – both available at the Chlamydomonas Center) were grown until mid-log phase in TAP medium with shaking at 100 rpm at 25°C in constant white light (54 µmol m^-2^s^-1^, provided by fluorescent bulbs). Intact chloroplasts were isolated as described (*95*) with slight modifications as follows: cells were harvested at 3,000 *g* for 4 min at 4°C and washed with 50 mM Hepes-KOH, pH 7.8. After centrifugation at 3,000 *g* for 5 min at 4°C, the cells were suspended in isolation buffer [50 mM Hepes-KOH, pH 7.8, 0.3 M sorbitol, 2 mM EDTA, 1 mM MgCl_2_, 0.1% (w/v) BSA, 0.5% (w/v) sodium ascorbate]. Just before cell breakage, cell suspensions were diluted to 0.5-3 mg chlorophyll / ml with isolation buffer and transferred to a 10 ml-Leuer-lock-syringe. The cells were broken by two passages through a 27-gauge needle at a flow rate of 0.1 ml / s. The suspensions were overlaid onto 45% / 80% Percoll step gradients [45% (v/v) or 80%(v/v) of Percoll, 50 mM Hepes-KOH, pH 6.8, 0.33 M sorbitol, 1 mM Na_4_P_2_O_7_, 2 mM EDTA, 1 mM MgCl_2_, 1 mM MnCl_2_, 0.3% (w/v) sodium ascorbate] and centrifuged in a swinging-bucket rotor at 4,200 *g* for 15 min at 4°C. Intact chloroplasts were collected from the 45% −80% interface, diluted with 5 volumes of HS buffer (50 mM Hepes-KOH, pH 7.8, 0.3 M sorbitol) followed by centrifugation at 1,000 *g* for 3 min at 4°C, and resuspended in HS buffer. After measuring chlorophyll concentration, chloroplasts were centrifuged at 1,600 *g* for 2 min at 4°C and stored either on ice or at −80°C.

### Preparation of urea-denatured pre-proteins

To express model pre-proteins in *E. coli*, cDNAs encoding pre-RbcS2, and pre-Fdx1 were obtained by RT-PCR from Chlamydomonas mRNA with the following primers: preRBCS2, CrRBCS2_F1_SpeI (5’-cctactagtGTCATTGCCAAGTCCTCCGTC-3’) and CrRBCS2_B1_BglII (5’-ccaagatctCACGGAGCGCTTGTTGGCGGG-3’); preCrFDX1, YACrFd1_F (SpeI) (5’-cttactagtATGGCCATGGCTATGCGCTCC-3’) and YACrFd1_R (Bgl II) (5’-cttagatctGTACAGGGCCTCCTCCTGGTG-3’). The cDNAs were cloned into pET24a-FLAG_3x_-TEV-Protein A-HIS_6x_ (*44*) to generate pET24-preCrRBCS2 and pET24-preCrFDX1. Model pre-proteins were expressed in *E. coli* BL21 (DE3) Star (Thermo Fisher Scientific) and purified on His•Bind resin (Novagen) in the presence of 8 M urea according to manufacturer’s instructions. The concentrations of pre-proteins were adjusted to 10-20 µM with 8 M urea buffer (8 M urea, 250 mM NaCl, 50 mM Tris-HCl, pH 7.5) and stored at −80°C until use.

### *In vitro* protein import experiments

*In vitro* import experiments with isolated intact chloroplasts prepared from Chlamydomonas strain CC-400 and purified pre-proteins were performed as described for *A. thaliana* (*44*) with the following minor modifications. Briefly, pre-proteins pre-RbcS2 and pre-Fdx1 with a C-terminal FLAG_3x_-TEV-Protein A-HIS_6x_ tag were microfuged for 10 - 20 min at 25°C to remove insoluble materials and denatured again with an equal volume of 8 M urea dilution buffer (8 M urea, 20 mM DTT, 10 mM Hepes-KOH, pH 7.8) immediately before use. Intact chloroplasts (1 - 2 mg chlorophyll) were incubated with denatured pre-proteins (100 - 200 nM) in 3 - 4 ml of HS buffer containing 0, 0.3, or 3 mM Mg-ATP, 5 mM MgCl_2_, 5 mM DTT) for 15 min at 25°C in the dark. After centrifugation at 1,600 *g* for 2 min at 4°C, chloroplasts were washed twice with HS buffer. Chloroplasts were resuspended with HS buffer with or without (for thermolysin treatment) 0.1 % protease inhibitor cocktail (for plant cells, Sigma, P-9959) and transferred to a new tube. Thermolysin treatment was carried out as previously described (*44*). For purification of translocation intermediates, chloroplasts were pelleted by centrifugation and snap-frozen in liquid nitrogen and stored at −80°C until use. To obtain soluble fractions containing stromal proteins, chloroplasts suspended in HS buffer containing 0.1% protease inhibitor cocktail were subjected to two freeze-thaw cycles, and the supernatant fraction was obtained after centrifugation at 21,500 *g* for 5 min at 4°C.

### Purification of translocation intermediates

Translocation intermediates after *in vitro* import experiments of model pre-proteins were purified as described for *A. thaliana* (*44*) with slight modifications. Stored chloroplasts were suspended in solubilization buffer (1% water-soluble digitonin, 50 mM Tris-HCl, pH 7.5, 10% [w/v] glycerol, 250 mM NaCl, 5 mM EDTA, 5 mM DTT, 0.5% protease inhibitor cocktail) to a final concentration of 2 mg chlorophyll / mL for 20 min with gentle rotation at 4°C. To remove insoluble materials, the chloroplast suspension was centrifuged at 21,500 *g* for 2 min at 4°C, and the supernatant was again ultracentrifuged with a Hitachi S100 AT5 angle rotor at 100,000 *g* for 5 min 4°C. The resulting supernatant (∼1 ml) was incubated with 20 µl of dimethyl pimelimidate (DMP) cross - linked IgG Sepharose 6 Fast Flow (GE healthcare) resin for 2 h with gentle rotation at 4°C. After washing 4 times with 0.7 ml of 0.2% digitonin-containing TGS buffer (50 mM Tris-HCl, pH 7.5, 10% [w/v] glycerol, 250 mM NaCl, 1 mM DTT), resins were further washed with the same buffer for 5 min with rotation at 4°C to remove non-specific proteins. The resins were transferred to a siliconized 0.5 ml-tube, and washed twice with 0.5 ml of 0.2% digitonin-containing TGS buffer. Bound translocation intermediates were eluted by cleavage with TEV protease in 100-120 µl of 0.2% digitonin containing TGS buffer for 1 h at 25 °C. To capture His-tagged TEV protease, 12 µl of complete His-Tag Purification Resin (Roche) was added and incubated for 15 min at 25°C. After centrifugation at 8,700 *g* for 1 min at 4°C, the supernatant was applied on a Micro Bio-spin column (Bio-Rad) to remove remaining resins, and the resulting flow-through was collected. For SDS-PAGE, the eluates were immediately denatured with sample buffer containing 16.7 mM Tris, pH 6.8, 50 mM (2-carboxyethyl) phosphine hydrochloride (TCEP-HCl) (Sigma, 66547) and 0.1% protease inhibitor cocktail at 37°C for 30 min. For mass-spec analysis, the eluates were stored on ice until use.

### Purification of the Chlamydomonas TicToc supercomplex containing FLAG3x-tagged Tic20

Chloroplasts (2-3 mg chlorophylls) isolated from CS1_FC1D12 (a CC-4533 strain transformed with a *TIC20-YFP-FLAG_3x_* nuclear transgene) (were solubilized with digitonin as described for the purification of translocation intermediates. Solubilized proteins were incubated with 15-20 µl of Anti-FLAG M2 Affinity Gel (Sigma) for 2 h with gentle rotation at 4°C. After similar extensive washing as described above, bound complexes were eluted with 100-120 µl of FLAG_3x_ peptide (100 µg / mL)(Sigma) in 0.2% digitonin containing TGS buffer for 40 min at 4°C.

### Production of Antisera

For expression of the Chlamydomonas Tic56 protein fragment in *E. coli*, the Cre17.g727100 coding sequence corresponding to amino acid residues 114-144 of Chlamydomonas Tic56 was synthesized; 5’-GGTGAACTGCGTCCGGTTCCGCGTAAAATTGTTCTGAGCCCGTATCAGTATGAGATGATTAACTATCAGCGTATGCTGATGCGCAAAAACATTTGGTATTATCGCGATCGTATGAATGTTCCGCGTGGTCCGTGTCCGCTGCATGTTGTTAAAGAAGCATGGGTTAGCGGTATTGTGGATGAAAATACCCTGTTTTGGGGTCATGGTCTGTATGATTGGCTGCCTGCAAAAAACATTAAACTGCTGCTGCCGATGGTTCGTACACCGGAAGTTCGTTT TGCAACCTGGATTAAACGTACCTTTAGCCTGAAACCGAGCCTGAATCGTATTCGTG AACAGCGTAAAGAACATCGTGATCCGCAAGAAGCAAGCCTGCAGGTTGAACTGAT GCGT-3’. The synthetic DNA fragment was cloned into the expression vector pGEM-EX1 (Promega) with a C-terminal HIS_6x_-tag. Polyclonal antiserum against Chlamydomonas Tic56 was produced by immunization of the purified Chlamydomonas Tic56 fragment as an antigen into a guinea pig. To produce the polyclonal antiserum against Chlamydomonas Tic214, a rabbit was immunized with a recombinant protein fragment corresponding to amino acid residue 628-737 of Chlamydomonas Tic214. This antigen was purified under denaturing conditions starting from BL21 *E.coli* cells transformed with pED3 (for detail, see construction of plasmids). A polyclonal antiserum against Chlamydomonas Tic20 was generated in collaboration with Yenzym, South San Francisco, upon rabbit immunization with the following peptide antigen: CRAEDAEKQDWKFGRNEG.

### Protein extraction and immunoblot analysis

Unless stated otherwise, total protein extraction and immunoblot analysis were performed as previously described (*70*). When a TCA protein extraction protocol was employed, the cell pellet was resuspended in 10% TCA in acetone plus 0.07% β-ME. To allow protein precipitation, the lysate was incubated at −20°C for at least 45 min and then subjected to centrifugation at 20,000g at 4°C for 15 min. The pellet was washed with cold acetone containing 0.07% β-ME at least twice. Finally, the pellet was dried with a speed vac and resuspended in a denaturing protein buffer containing 50 mM Tris-HCl pH 6.8, 300 mM NaCl, 2% SDS, and 10 mM EDTA. Prior to immunoblot analysis, the protein content of each sample was measured by BCA assay to ensure equal loading. Transfer to PVDF membranes was carried out with (for Tic214 and others) or without (for Tic56) 0.01% SDS in transfer buffer at 60 V for 2 h or 20 V overnight on ice. Proteins were detected by the ECL Prime Western Blotting System (GE healthcare) or Clarity ECL Western Substrate (Bio-Rad) and exposed to X-ray films (Super RX, Fuji film).

### LC-MS/MS analysis of translocation intermediates

After alkylation with iodoacetamide, purified proteins were digested with trypsin in solution. LC-MS/MS analysis was performed by UltiMate 3000 Nano LC systems coupled to Q-Exactive hybrid quadrupole-Orbitrap mass spectrometer (Thermo Fisher Scientific). Peptides and proteins were identified by Mascot v2.3 (Matrix Science, London) searched against the Uniprot Chlamydomonas dataset and the *Creinhardtii*_281_v5.5.protein datasets.

### LC-MS/MS analysis of total protein extracts upon Tic214 depletion and proteasome inhibition

*<u>Cell pellets preparation</u>*: at the beginning of the experiment, 20-ml aliquots of cell culture in the late exponential phase (at a cell density of about 7*10 ^6^ cells / ml) were inoculated and diluted ten times either in regular TAP or TAP freshly supplied with 200 µM Thiamine and 20 µg / ml B12. Thereafter, to keep cell growth in the exponential phase, all cultures were diluted every 24 h to a final cell concentration of about 7*10^5^ cells/ml. At each harvesting time point (2, 4, and 6 days), about 4*10^8^ cells were pelleted at 1000 g for 5 min at RT. After being resuspended in 25 ml of the same type of growth medium (i.e., / + Vit), they were incubated for 3 h in the presence of 30 µM MG132 (Sigma Aldrich #M7449). Then, cells were pelleted again, frozen in liquid nitrogen, and stored at −80°C until use. *<u>Cell pellets processing</u>*: the weight of each frozen cell pellet was measured and resuspended in 5 volumes of lysis buffer containing 100 mM Tris-HCl pH8, 600 mM NaCl, 4% SDS, 20 mM EDTA and freshly supplemented with MS-SAFE Protease and Phosphatase Inhibitor Cocktail (Sigma Aldrich # MSSAFE) (e.g., 100 mg frozen pellet = 500 µl lysis buffer). Cells were disrupted by constant agitation in this buffer for 30 min at 4°C. Then, the protein mixture was further denatured for 30 min at RT and centrifuged at 21000 g for 30 min at 4°C to remove cellular debris. The supernatant (i.e., total protein) was transferred in a clean Eppendorf, and a 5-µl aliquot of this clear lysate was used to determine protein concentration by BCA assay (REF. Perlaza et al. 2019). *<u>Sample preparation for Mass-spec analysis</u>*: for each time point, 60 µg of total protein extracts were mixed with Laemmli sample buffer (Biorad #1610747), freshly supplemented with 2-mercaptoethanol (Biorad #1610710), heated for 30 min at 37°C and loaded into a polyacrylamide gel (any kD™ precast protein gel, Biorad #4569034). To avoid cross-contamination, one empty well was left between the different protein samples. Proteins were allowed to electrophorese into the gel for 15 min at 100 V, visualized by Colloidal Coomassie staining (REF. https://www.ncbi.nlm.nih.gov/pmc/articles/PMC3149902/), excised as a single gel band of 0.5 × 1.5 cm, transferred in a clean Eppendorf tube a 1% acetic acid solution, and sent out to the Stanford University Mass Spec Core. *<u>Mass spec analysis</u>*: the identification of peptides and proteins by tandem mass spectrometry was carried out by the Stanford University Mass Spec Core using the Byonic software package. Output data were organized in two different types of Excel spreadsheets, one for proteins, and one for peptide-spectrum matches (PSMs) and were summarized in a heatmap (*SI Appendix*, Table S10). *<u>Data mining</u>*: Only proteins for which at least 10 MS-peptides could be identified in one of the six conditions were taken into consideration for further analysis. Their localization was predicted using the Predalgo software (*65*). In 80% of the cases (345/427), the chloroplast localization was confirmed by another prediction software, ChloroP (*96*). Only proteins for which sequences derived from their predicted cTP could be detected upon Tic214 were annotated as potential TIC clients. This information is available in *SI Appendix*, Table S11. The PhytoMine interface (https://phytozome.jgi.doe.gov/phytomine) was used to identify potential *A. thaliana* orthologs of these proteins listed in *SI Appendix*, Table S12. To assess whether Tic214 knockdown affects chloroplast stress-responsive proteins encoded by nuclear genes and *MARS1*-dependent genes, we added +1 to all spectral counts, and determined the protein fold-change for each timepoint as log_2_ (spectral counts Tic214 OFF / spectral counts Tic214 ON). A protein was considered affected by Tic214 knockdown when its fold change was = or > 2 at least in one timepoint (768 proteins). This information is available in *SI Appendix*, Table S13. Lists of chloroplast stress-responsive genes and *MARS1*-dependent genes were extracted from (*75*). All calculations were done in R (R project v3.5.1) (www.R-project.org) using a combination of the packages stringr (https://CRAN.R-project.org/package=stringr.), dplyr (https://CRAN.R-project.org/package=dplyr.), gplots (.https://CRAN.R-project.org/package=gplots.) and custom scripts.

### Miscellaneous

Isolation of RNA and DNA from Chlamydomonas strains and PCRs on genomic DNA to test chloroplast genome homoplasmicity were performed as previously described (*70*). RT-PCRs were performed as previously described (*44*). Mass-spec compatible silver staining was carried out as previously described (*97*). 2D Blue native/SDS-PAGE was performed as previously described (*98*). Electron microscopy imaging was carried out as previously described (*71*). Co-immunoprecipitation studies shown in Table 1 and *SI Appendix*, Fig. S3 and Tables S6 and S7 were performed as previously described (*61*).

## Authors Contributions

S.R., J.D.R, M.N., and P.W. conceived and planned experiments; S.R., Y.A., K.T., and M.C. carried out experiments; L.C.M.M., M.C.J., M.R., and E.D. help to generate critical reagents; S.R., Y.A., P.S., D.S., M.B., K.T., L.M., M.H., M.N., J.D.R., and P.W. analyzed data; S.R. and P.W. wrote the manuscript with inputs from all authors. All authors read and approved the manuscript.

## Acknowledgments

We thank Christopher M. Adams and Ryan Leib from the Stanford University Mass Spectrometry Platform for carrying out mass-spectrometry analysis, Lorenzo Costantino, Elif Karagoz, and Jan Schuller for useful feedback on the manuscript. This project was supported by an EMBO long term-fellowship (ALTF 563-2013) and a Swiss National Science Foundation Advanced PostDoc Mobility Fellowship (P2GEP3_148531) awarded to SR, a Belgian-American Educational Foundation fellowship to M.B., a Humboldt Research Fellowship awarded to L.M., a Deutsche Forschungsgemeinschaft grant (HI 739/9.1 and HI 739/9.2) awarded to M.H., two grants from Ministry of Education Culture, Sports, Science and Technology (17H05668 and 17H05725) awarded to MN, a cooperative agreement of the US Department of Energy Office of Science, Office of Biological and Environmental Research program (DE-FC02-02ER63421) awarded to Sabeeha Merchant and Todd Yeates (UCLA), and a grant from the Swiss National Foundation (31003A_133089/1) awarded to JDR. PW is an Investigator of the Howard Hughes Medical Institute (HHMI826735-0012) and is supported by the National Institute of Health (R01GM032384).

---

**Fig. S1.**
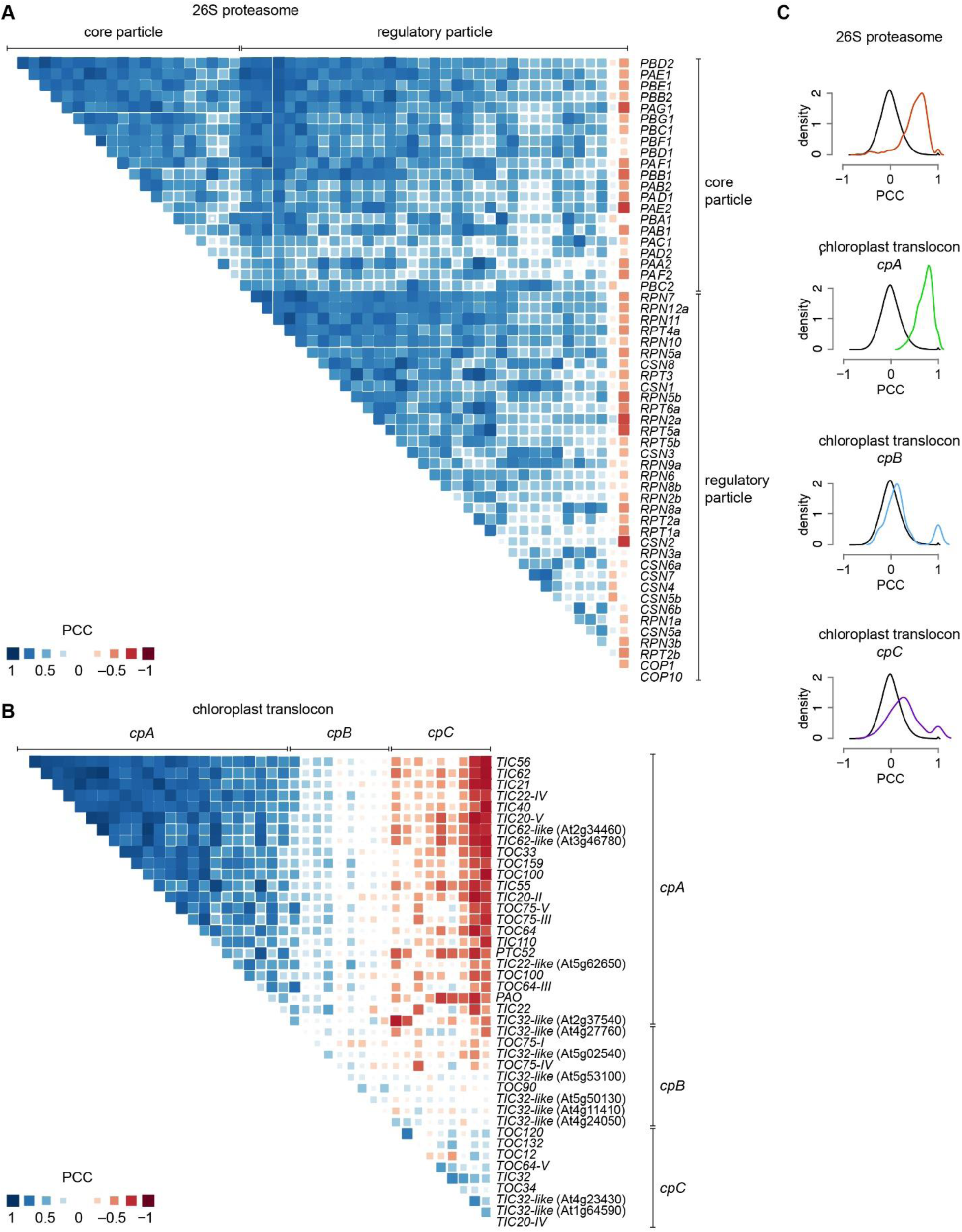
Co-expression patterns of Arabidopsis genes encoding components of the plastid translocon. (*A*) Correlation matrix for Arabidopsis genes encoding subunits of the 26S proteasome (listed in *SI Appendix*, Table S5). (*B*) Correlation matrix for Arabidopsis genes encoding components of the chloroplast translocon (listed in *SI Appendix*, Table S5), clustered in three sub-groups: *cpA*, *cpB,* and *cpC*. (*C*) PCC distribution for all gene pairs encoding core and regulatory particles of the 26S proteasome (red line), for all gene pairs encoding chloroplast translocon components as a function of their subgroup — *cpA* (green line), c*pB* (blue line) and *cpC* (purple line) — and for all gene pairs in the genome (black line) used as a negative control. Statistical significance of all distributions was tested by Kolmogorov-Smirnov test, comparing each sets of PCC values with a randomly generated normal distribution of equal element number. The results of this analysis are available in *SI Appendix*, Table S4.

**Fig. S2.**
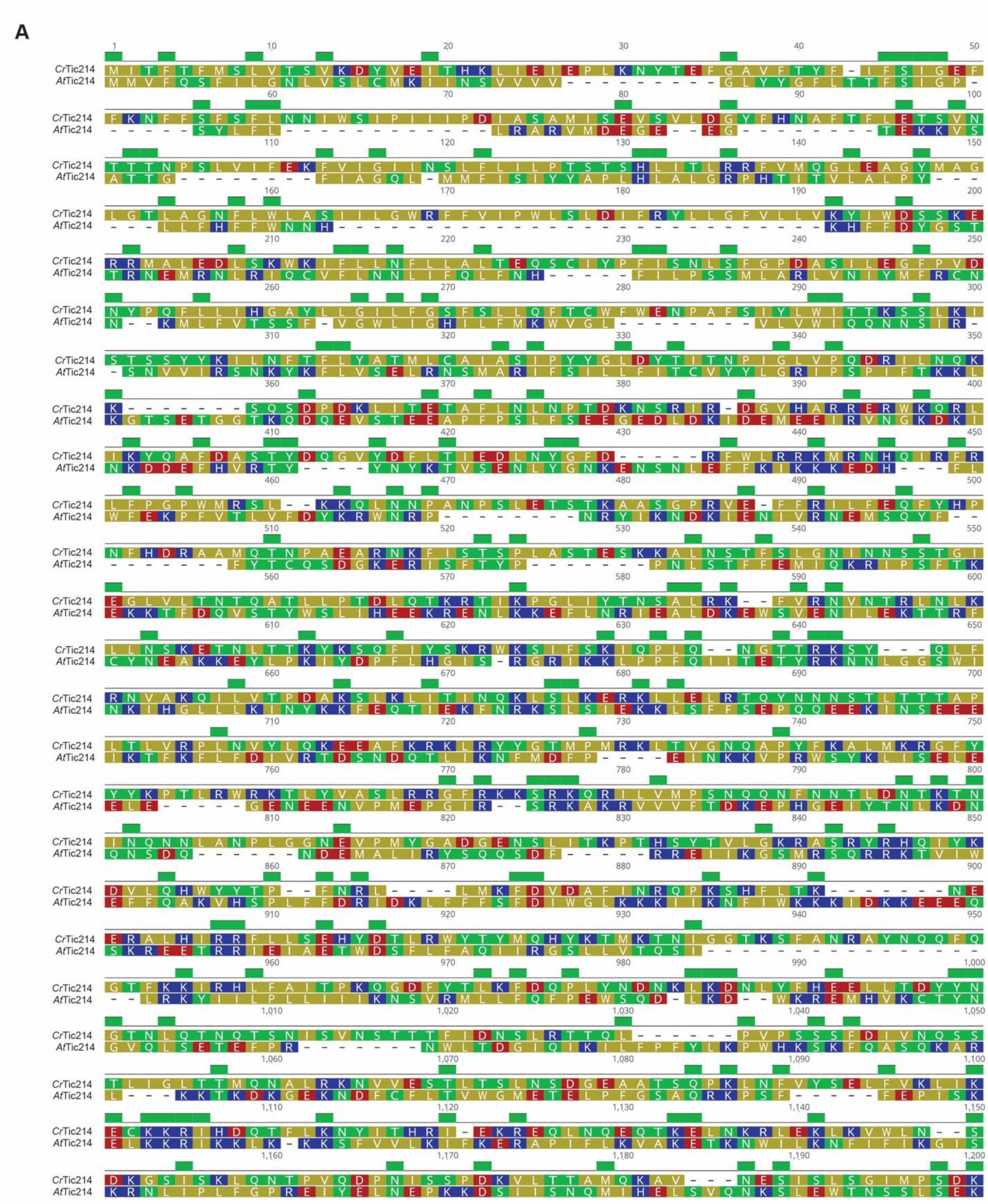
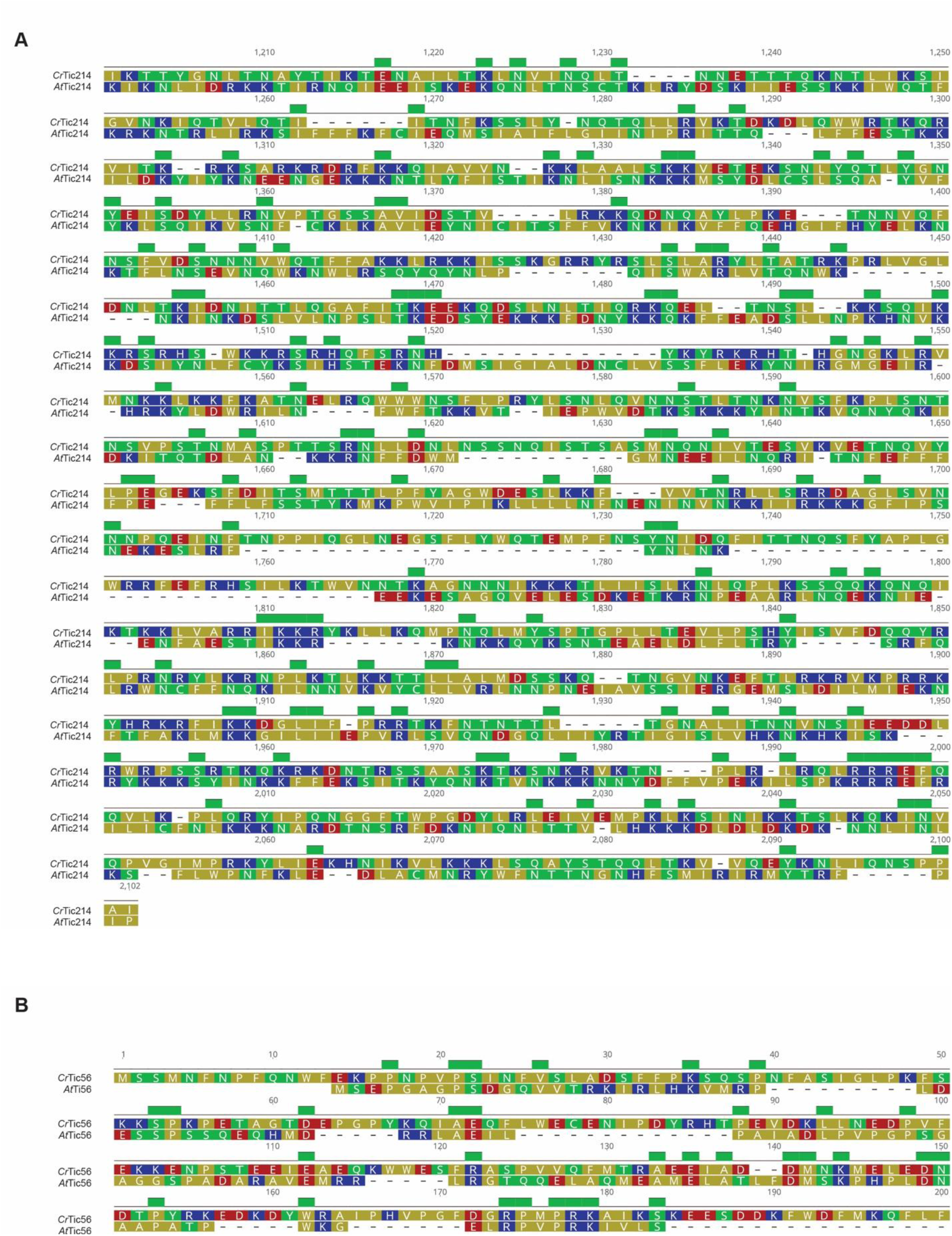
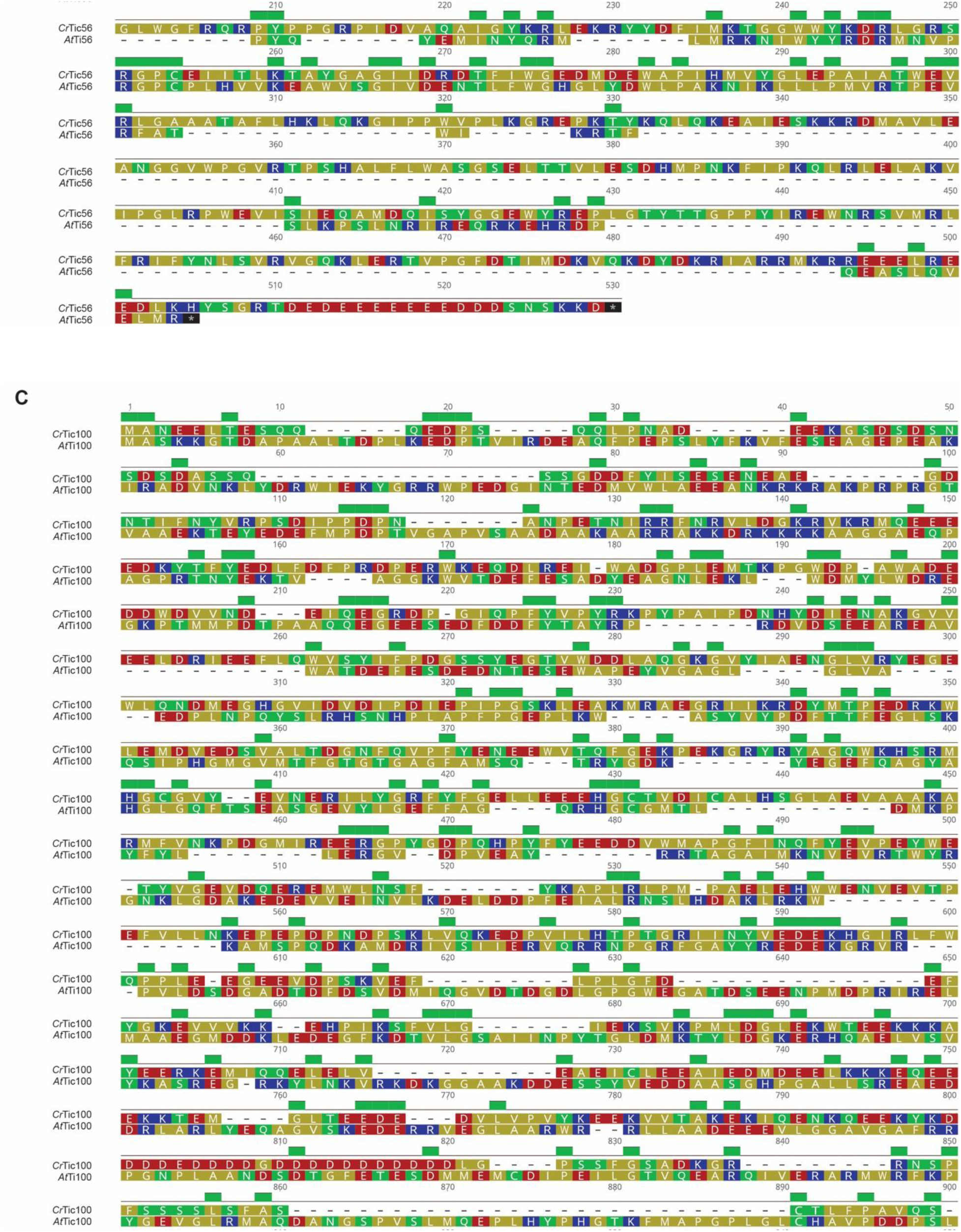
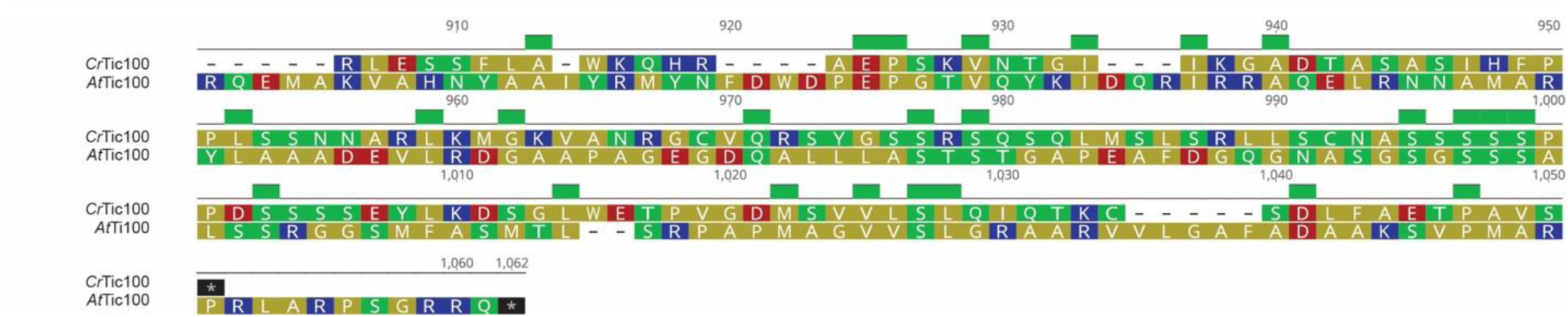
Sequence comparison of TIC components of Chlamydomonas and Arabidopsis. All protein alignments were performed using Geneious 11.1.5 (alignment method: MUSCLE, Topology: LINEAR). The amino acids were colored according to their polarity as follows: Yellow = Non-polar (G, A, V, L, I, F, W, M, P), Green = Polar, uncharged (S, T, C, Y, N, Q), Red = Polar, acidic (D, E), Blue = Polar, basic (K, R, H). (*A*) Tic214 sequence comparison of Arabidopsis and Chlamydomonas (*orf1995*/*ycf1*). Identity = 321/2102 (15%), Positives = 715/2102 (34%), Gaps = 423/2102 (20%). (*B*) Tic56 sequence comparison of Arabidopsis and Chlamydomonas (At5g01590/Cre17.g727100). Identity = 71/530 (13%), Positives = 120/530 (22%), Gaps = 287/530 (54%). (*C*) Tic100 sequence comparison of Arabidopsis and Chlamydomonas (At5g22640/Cre06.g300550). Identity = 179/1062 (16%), Positives = 321/1062 (30%), Gaps = 296/1062 (27%).

**Fig. S3:**
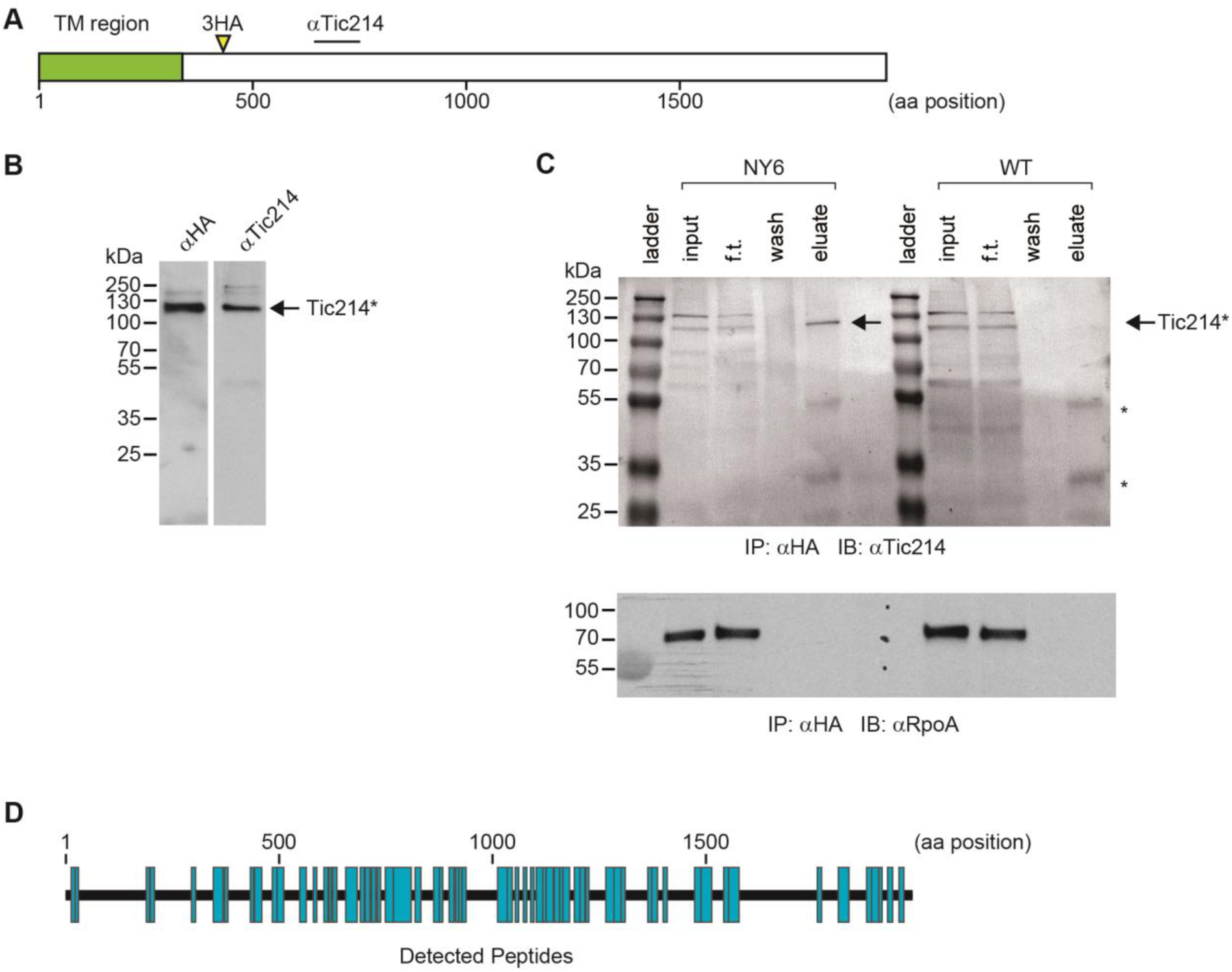
Characterization of Tic214. (*A*) Organization of Tic214 with 8 N-terminal transmembrane domains and the remaining basic region. The region used for purifying a recombinant protein and producing antibodies (αTic214) and the insertion site of the HA-epitope after T402 (3XHA), are indicated. (*B*) Immunoblot analysis of total protein of NY6 cells containing HA-tagged Tic214 with anti-HA and anti-Tic214 antibodies. Tic214* denotes the 110 kDa protein band detected by the Tic214 antibody. (*C*) Co-immunoprecipitation of total protein from NY6 containing HA-tagged Tic214. Cellular extracts from NY6 and wild type (WT) were incubated with an affinity matrix containing HA antibodies. After extensive washing of the matrix, bound proteins were eluted with SDS buffer and immunoblotted with HA antiserum. The black arrows highlight the position of the 110 kDa protein band detected by the Tic214 antibody in the immunoblot. (*D*) The entire *tic214* mRNA is translated as a protein of 232 kDa. After immunoprecipitation with anti-HA antibodies, proteins were digested with trypsin and analyzed by mass spectrometry. The distribution of the identified peptides of Tic214 (indicated in blue) over the entire Tic214 sequence (between D15 and K1979) is drawn to scale.

**Fig. S4.**
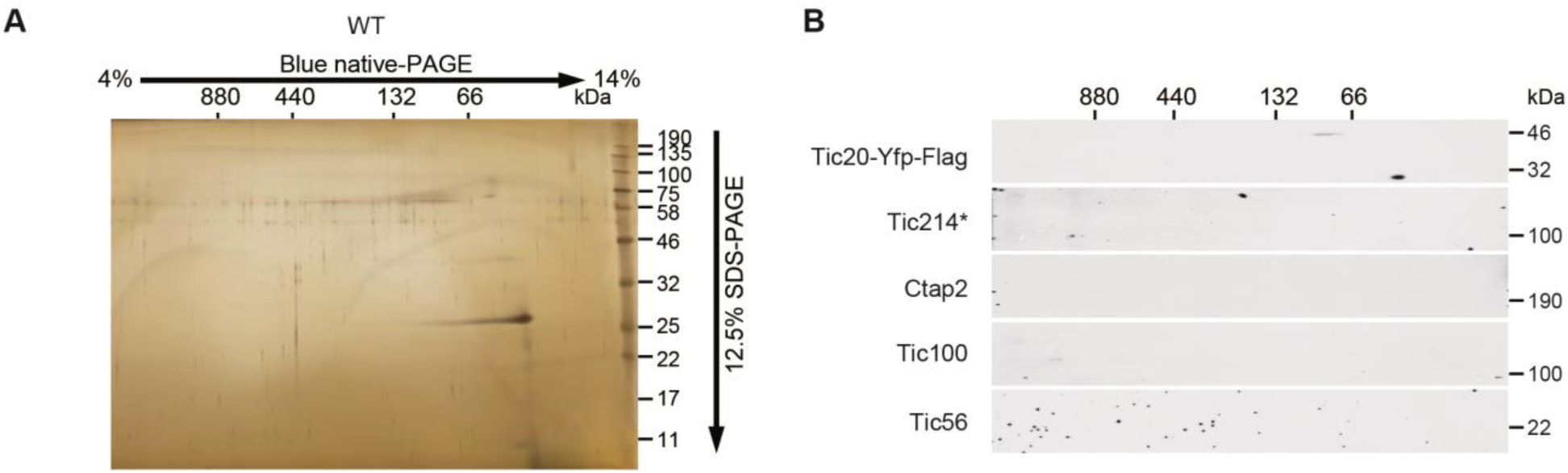
2D blue native/SDS-PAGE separation of a mock purified sample. (*A*) A mock purified sample with untagged Tic20 was analyzed as in Fig. 4A. (*B*) A mock purified sample with untagged Tic20 was analyzed as in Fig. 4B.

**Fig. S5.**
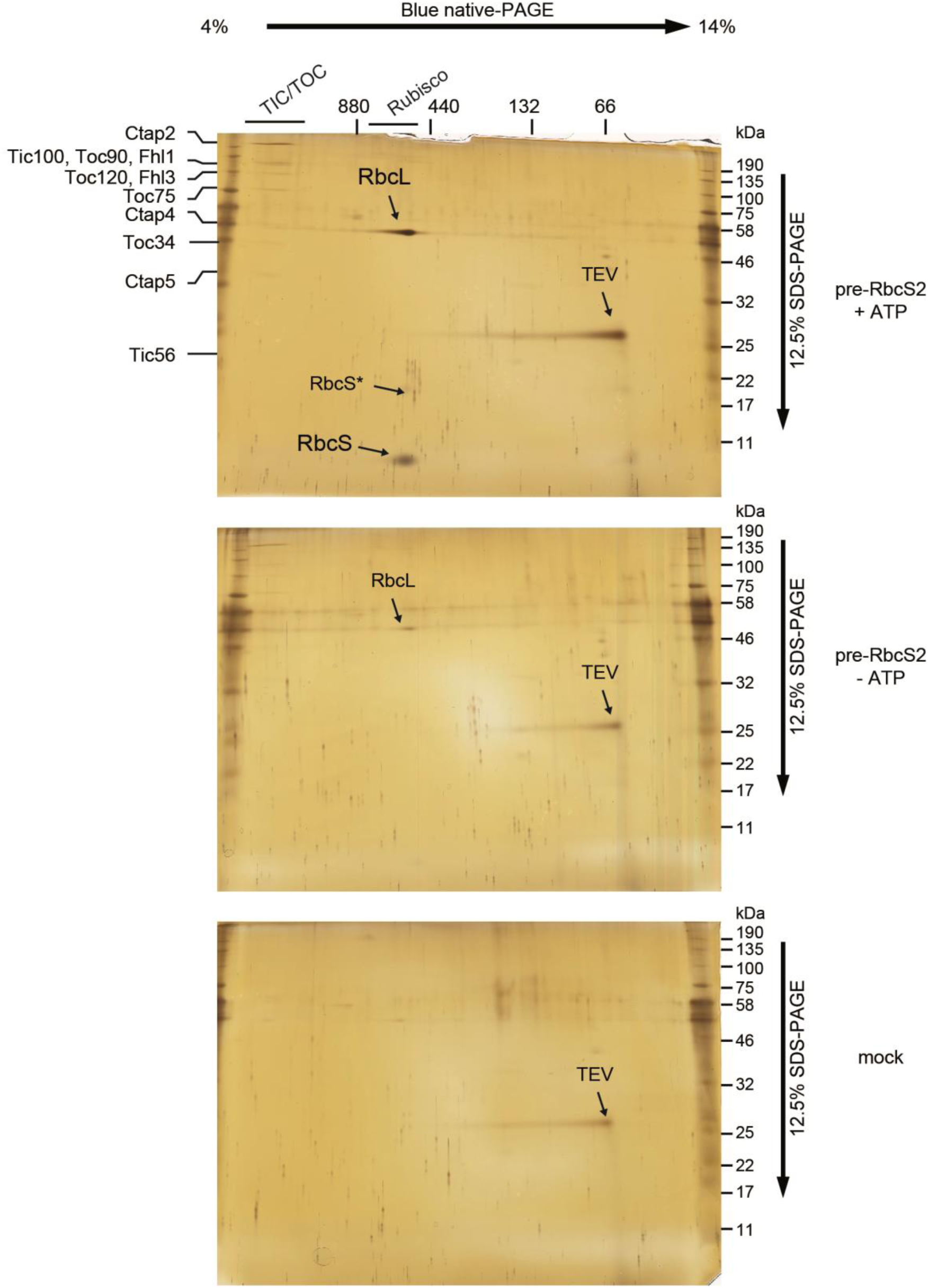
2D blue native/SDS-PAGE analysis of purified translocation intermediates. The pre-RbcS2 was used for in vitro import experiments with Chlamydomonas chloroplasts in the presence or absence of ATP. Translocation intermediates were purified and analyzed by 2D blue native/SDS-PAGE separation followed by silver staining. Mock-purified samples prepared from the same amounts of chloroplasts without the addition of pre-proteins were also analyzed. Bands containing proteins identified by mass spectrometry are labeled.

**Fig. S6.**
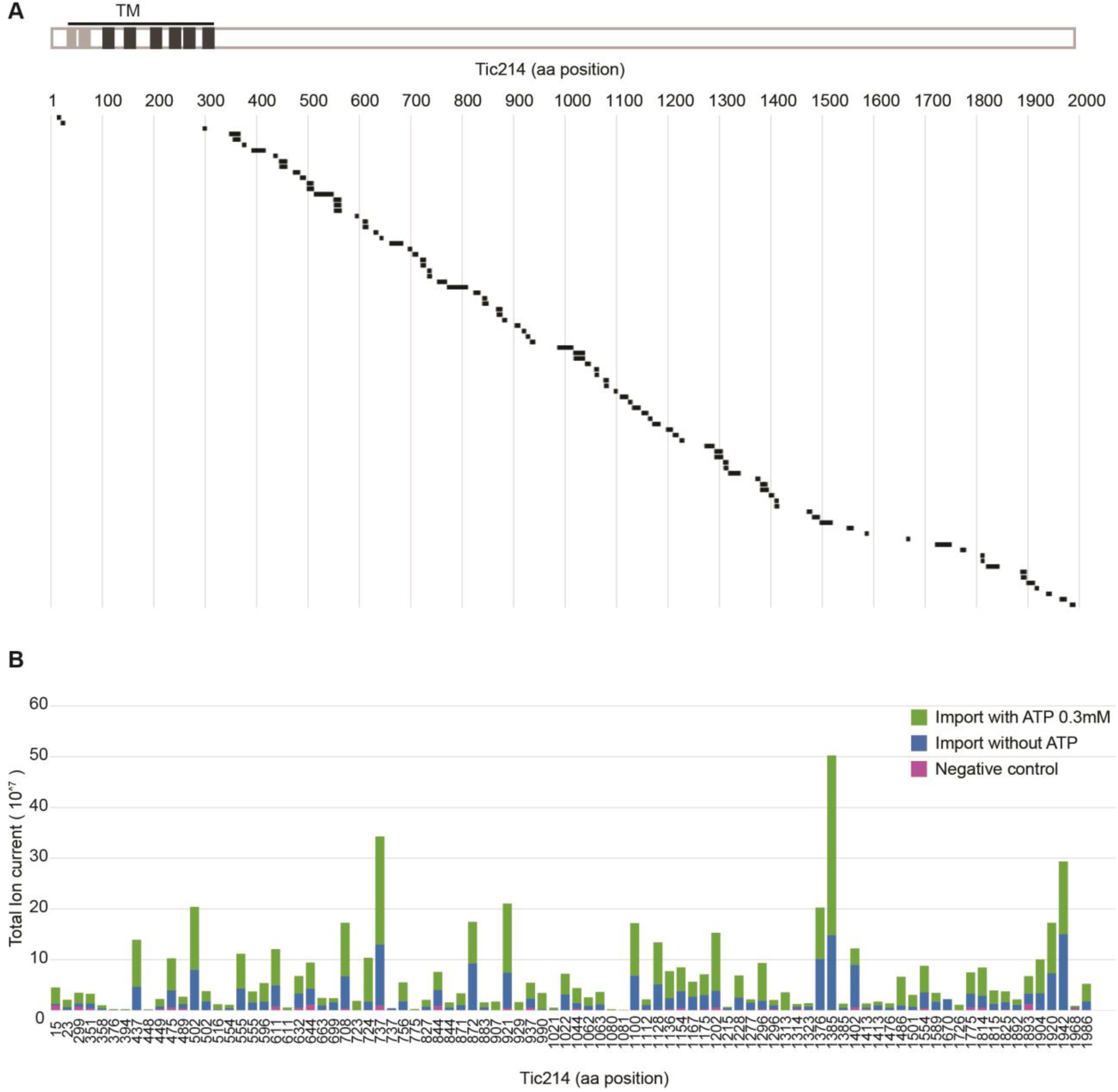
Distribution of Tic214-derived peptides identified by LC-MS/MS analysis of the translocation intermediates. (*A*) (*upper panel*) Strongly and weakly predicted transmembrane domains in Tic214 protein are shown in dark and light grey, respectively. (*lower panel*) Tic214-derived peptides identified upon mass spec analysis of translocation intermediates are indicated as dark bars along the length of the protein. (*B*) ATP-dependent association of Tic214 with translocating pre-proteins is shown as measured by total ion current detected for Tic214-derived peptides. The numbers of the starting amino acids of the identified peptides in full-length Tic214 are shown below the horizontal axis. In the negative control, no pre-protein was added to the chloroplasts.

**Fig. S7.**
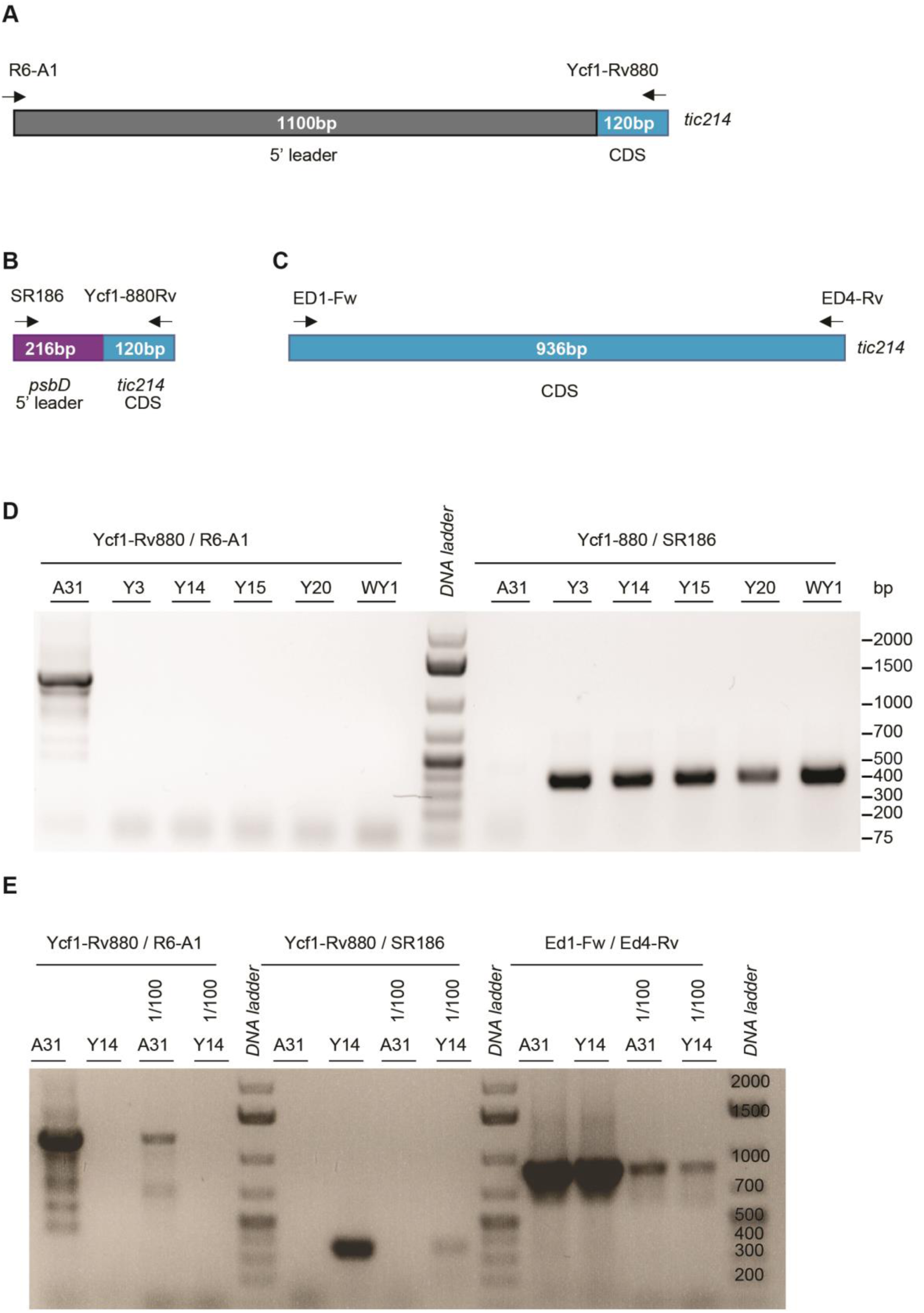
Homoplasmicity of the Y14 strain. (*A*) Location of primers used for PCR on authentic *tic214 (orf1995)* leader and coding sequence (CDS). (*B*) Location of primers used for PCR on chimeric *psbD*:*tic214* gene. (*C*) Location of primers used for PCR on *tic214* CDS. (*D*) Y strains were obtained by transformation of A31 with the pRAM73.19 plasmid containing the *psbD* 5’ leader fused to the CDS of *tic214* and the *aadA* spectinomycin resistance cassette. The WY1 strain was obtained in the same way except that the wild-type cell line was used for transformation. The authentic *tic214* locus was examined by PCR using the primers shown in panel A, while the chimeric *psbD*5’UTR:*tic214* was analyzed by PCR using primers shown in panel B. (*E*) The same PCR reactions shown in panel D were repeated for A31 and Y14. In this case, a 1/100 dilution of the genomic DNA was also tested to make sure that at least one gene copy per chloroplast is detectable as there are ∼80 copies of chloroplast DNA molecules per chloroplast in Chlamydomonas. Primers shown in panel C (spanning a region of *tic214* CDS) were used as loading control.

**Fig. S8.**
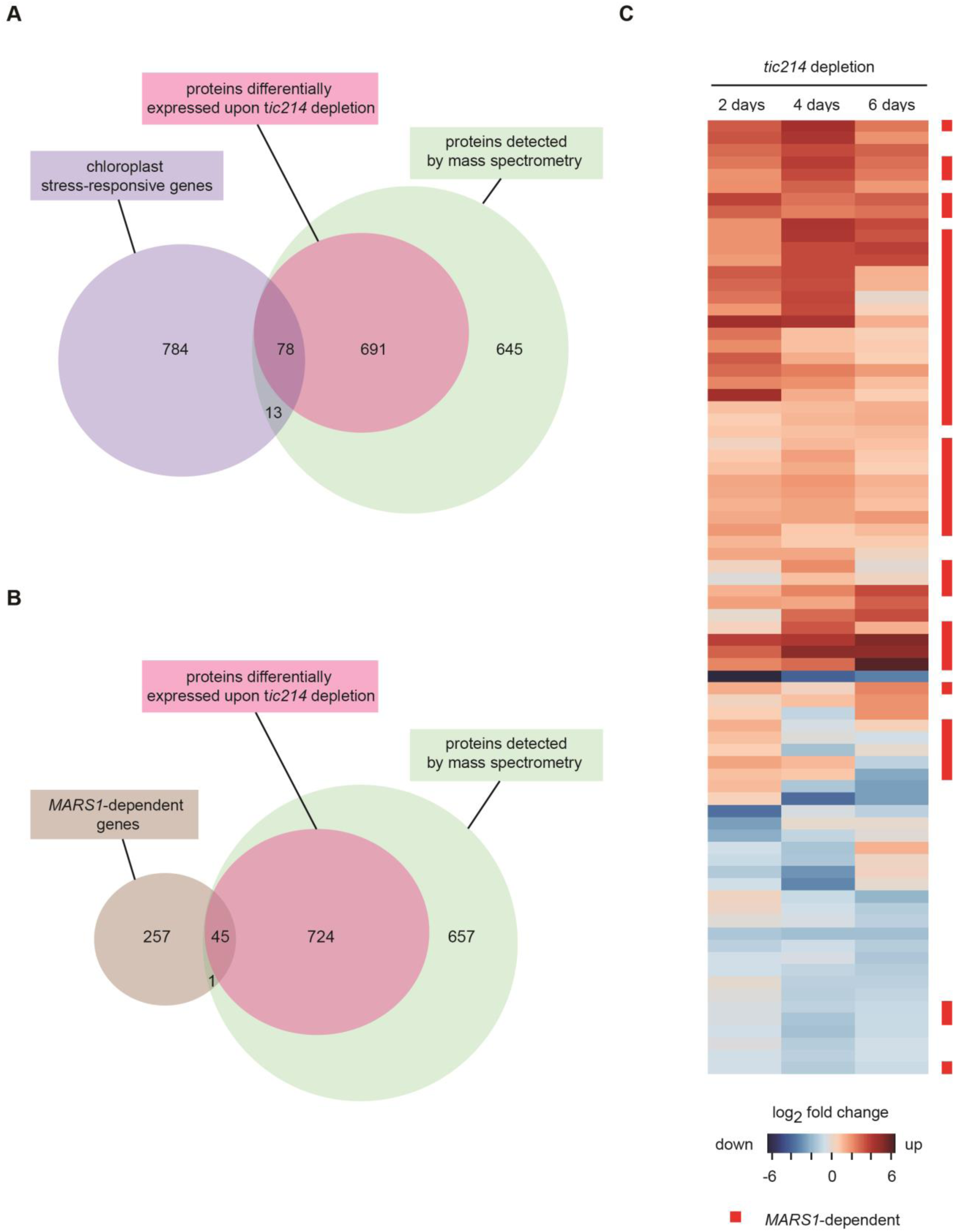
Chloroplast-stress responsive proteins differentially expressed upon Tic214 depletion. (*A*) Venn diagram highlighting the number of chloroplast stress-responsive proteins that could be detected by mass spectrometry (n = 91). About 80% (n = 78) were also differentially expressed upon *tic214* depletion (Protein IDs are listed in *SI Appendix*, Table S13). Chloroplast stress-responsive proteins are defined as proteins encoded by nuclear genes differentially expressed upon down-regulation of the chloroplast Clp protease and in response to excessive light in Chlamydomonas cells (*75*). (*B*) Venn diagram highlighting the number of proteins encoded by *MARS1*-dependent genes and detected by mass spectrometry (n = 46). Over 95% (n = 45) were differentially accumulated upon *tic214* depletion (Protein IDs are listed in *SI Appendix*, Table S13). *MARS1* encodes a critical component of the chloroplast unfolded protein response (*75*). (*C*) Heatmap showing the proteins encoded by chloroplast stress-responsive genes and differentially accumulated upon *tic214* depletion. The red squares on the side indicate those proteins encoded by *MARS1*-dependent genes.

## List and titles of Supplementary Tables

**Table S1.** Arabidopsis components of the TOC complex and their putative Chlamydomonas orthologs from BLAST analysis.

| Arabidopsis |  |  | Chlamydomonas |  |  | BLAST |  |
| --- | --- | --- | --- | --- | --- | --- | --- |
| Gene ID | Protein name | Protein length (aa) | Gene ID | Protein name | Protein length (aa) | Score | p-value |
| At5g05000 | Toc33 | 313 | Cre06.g252200 | Toc33/34 | 397 | 150.2 | 1.60E-41 |
|  |  |  | Cre17.g734300 | Toc90 | 967 | 62.8 | 7.60E-11 |
|  |  |  | Cre17.g707500 | Toc120 | 1080 | 58.2 | 2.70E-09 |
| At1g02280 | Toc34 | 297 | Cre06.g252200 | Toc33/34 | 397 | 145.6 | 1.50E-39 |
|  |  |  | Cre17.g734300 | Toc90 | 967 | 67 | 3.60E-12 |
|  |  |  | Cre17.g707500 | Toc120 | 1080 | 55.1 | 3.60E-08 |
| At1g08980 | Toc64 | 425 | Cre11.g467630 | Toc64 | 540 | 318.9 | 3.50E-103 |
| At3g17970 |  | 589 | Cre11.g467630 |  |  | 250.8 | 3.40E-75 |
| At5g09420 |  | 603 | Cre11.g467630 |  |  | 226.1 | 6.10E-66 |
| At3g46740 | Toc75-III | 818 | Cre03.g175200 | Toc75 | 798 | 303.9 | 8.80E-90 |
| At4g09080 | Toc75-IV | 396 | Cre03.g175200 |  |  | 222.2 | 2.30E-64 |
| At5g19620 | Toc75-V | 732 | Cre09.g388097 | Oep80 | 651 | 443 | 6.40E-145 |
| At5g20300 | Toc90 | 793 | Cre17.g734300 | Toc90 | 967 | 163.3 | 2.60E-41 |
| At4g15810 | Toc100 | 918 | Cre17.g734300 | Toc90 | 967 | 49.7 | 9.70E-06 |
| At3g16620 | Toc120 | 1089 | Cre17.g707500 | Toc120 | 1080 | 172.6 | 1.50E-43 |
| At2g16640 | Toc132 | 1206 | Cre17.g707500 | Toc120 | 1080 | 171 | 6.00E-43 |
| At4g02510 | Toc159 | 1503 | Cre17.g707500 | Toc120 | 1080 | 153.7 | 2.00E-37 |

(Supplemental data for Fig. 1)

**Table S2.**
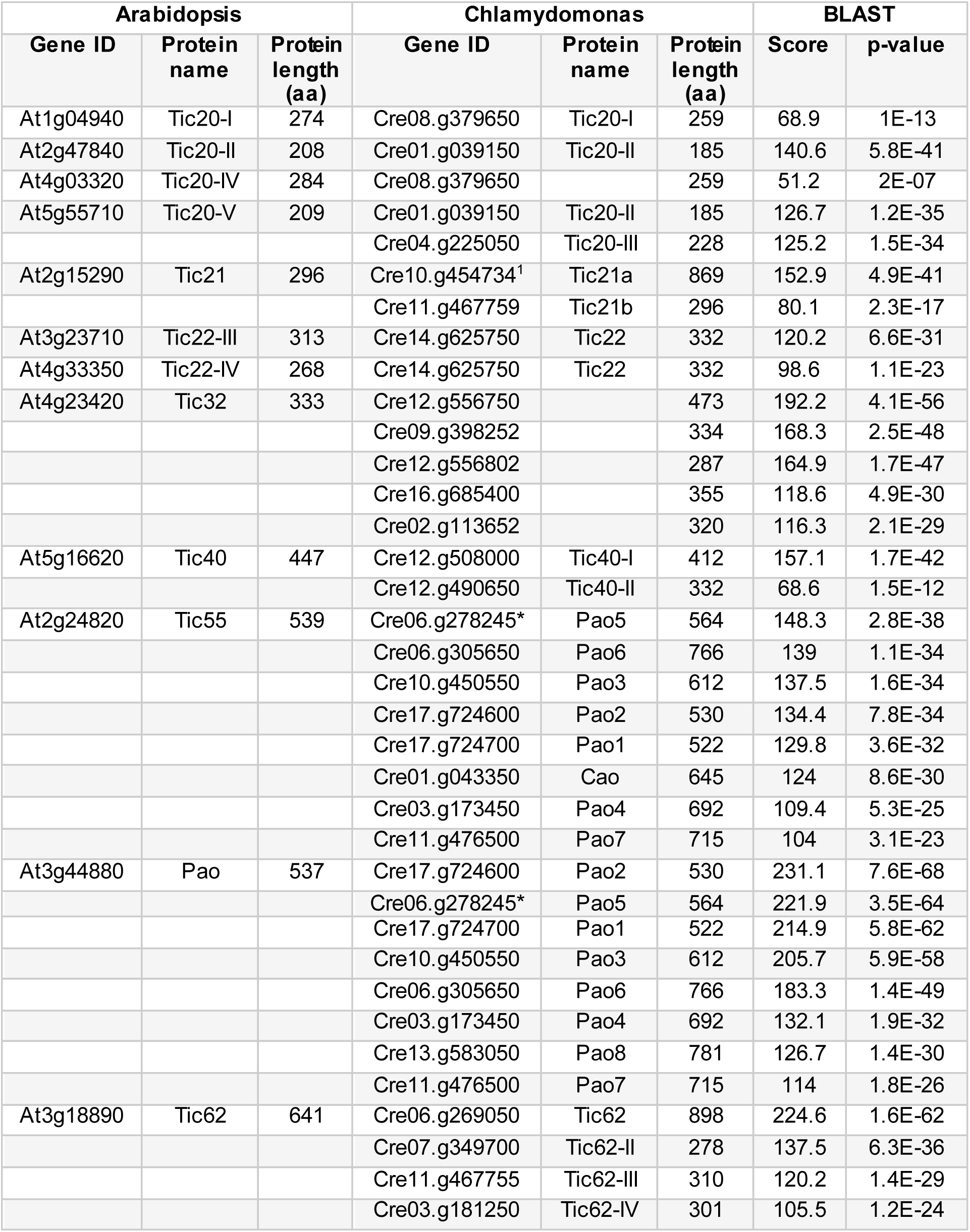
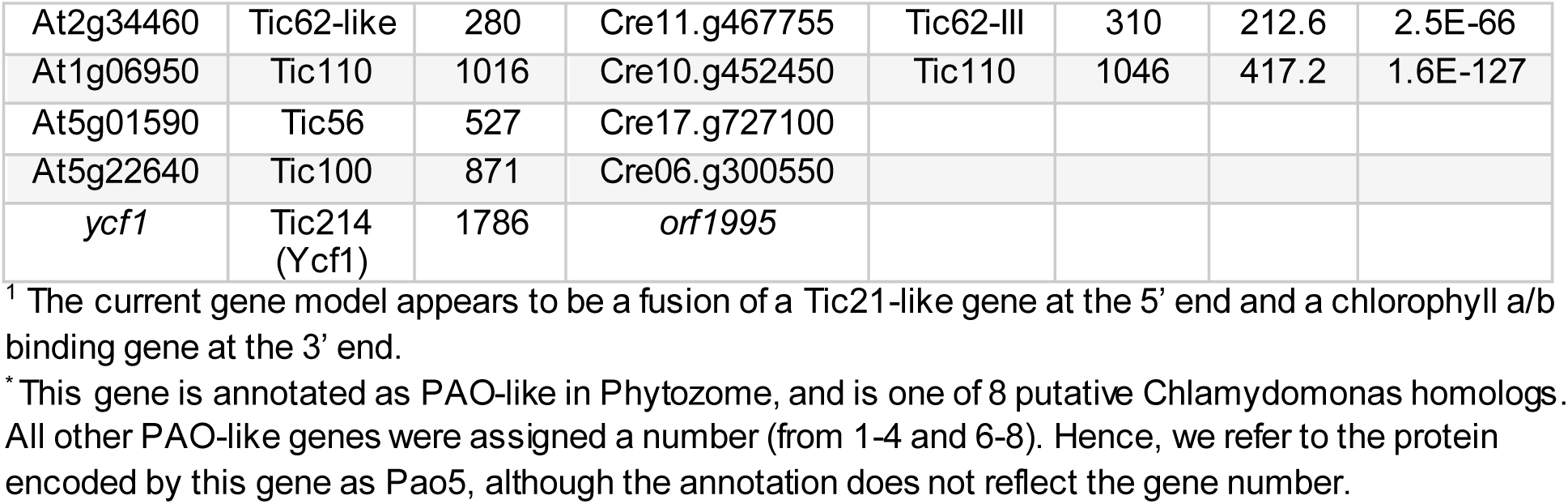
Proposed Arabidopsis components of the TIC complex and their putative Chlamydomonas orthologs from BLAST analysis.

(Supplemental data for Fig. 1 and Fig. 2)

Table S3. *(Excel file)* List of genes of the components of the proteasome and chloroplast translocons of Chlamydomonas shown in Fig. 1 *A* and *B*, respectively. (Supplemental data for Fig. 1)

Table S4. *(Excel file)* Statistical table related to data shown in Fig. 1, Fig. 2 and Fig. S1

Table S5. *(Excel file)* List of genes of the components of the proteasome and chloroplast translocons of Arabidopsis shown in Fig. S1 *A* and *B*, respectively.

Table S6. *(Excel file)* LC-MS/MS identification of proteins co-immunoprecipitated with Chlamydomonas Tic20.

(Supplemental data for Fig. 2 and Table1)

Table S7. *(Excel file)* LC-MS/MS identification of proteins co-immunoprecipitated with Chlamydomonas Tic214.

(Supplemental data for Fig. 2 and Table1)

**Table S8.** Genes co-expressed with Chlamydomonas *TIC20*.

| Gene ID | Mutual rank | Protein Name (Annotation) |
| --- | --- | --- |
| Cre06.g300550<sup>a,b</sup> | 8 | Tic100 |
| Cre16.g696000<sup>a,b</sup> | 3 | Ctap2 |
| Cre17.g727100<sup>b</sup> | 6 | Tic56 |
| Cre07.g352350<sup>b</sup> | 10 | Fhl3 |
| Cre03.g201100<sup>b</sup> | 6 | Fhl1 |
| Cre17.g739752<sup>b</sup> | 47 | Ctap1 |
| Cre12.g532100<sup>b</sup> | 9 | Ctap3 |
| Cre17.g722750<sup>b</sup> | 19 | Ctap4 |
| Cre03.g164700<sup>b</sup> | 22 | Ctap5 |
| Cre08.g378750<sup>b</sup> | 21 | Ctap7 |
| Cre03.g175200<sup>a,b</sup> | 5 | Toc75 |
| Cre17.g734300<sup>a,b</sup> | 4 | Toc90 |
| Cre08.g379650<sup>a,b</sup> | 1 | Tic20 |
| Cre06.g252200<sup>a,b</sup> | 21 | Toc34 |
| Cre02.g080250 | 4 | Ylmg1 (YGGT family) |
| Cre13.g573900 | 6 | Nss3 (Sodium/solute transporter) |
| Cre03.g201750 | 6 | not annotated |
| Cre12.g527550 | 8 | not annotated |
| Cre10.g451900 | 10 | Ths1 (Threonine synthase) |
| Cre13.g604650 | 11 | Metallopeptidase family M24 |
| Cre16.g683081 | 11 | Sec-C motif domain-containing protein |
| Cre12.g497850 | 12 | not annotated |
| Cre08.g365600 | 12 | Hydroxymethylpyrimidine kinase |
| Cre09.g386200 | 12 | Opr36 |
| Cre02.g142246 | 13 | not annotated |
| Cre04.g216950 | 15 | KasIII (Beta-ketoacyl-[acyl-carrier-protein] synthase III) |
| Cre03.g160500 | 16 | Tsk1 (Lysine-tRNA ligase) |
| Cre16.g659850 | 16 | Cgl37 (Shikimate kinase-related protein) |
| Cre16.g663150 | 17 | Thiosulfate sulfurtransferase |
| Cre03.g171100 | 17 | not annotated |
| Cre01.g052250 | 17 | Trx1 (Thioredoxin x) |
| Cre04.g214501 | 18 | Pnp1 (Polynucleotide phosphorylase) |
| Cre16.g691000 | 19 | Efp1 (Organellar elongation factor P) |
| Cre12.g552850 | 19 | Cgl77 |
| Cre09.g403145 | 20 | not annotated |
| Cre17.g747297 | 20 | peptidyl-tRNA hydrolase |
| Cre14.g629650 | 21 | Nik1 (Nickel transporter) |
| Cre05.g240850 | 23 | ThiC (Hydroxymethylpyrimidine phosphate synthase) |
| Cre09.g405150 | 24 | Pus1 (tRNA-pseudouridine synthase) |

<sup>a</sup> gene is part of the plastid translocon based on our BLAST analysis (SI Appendix, Tables S1 and S2)<sup>b</sup> encoded protein was identified during the co-immunoprecipitation studies presented in this manuscript.

(Supplemental data for Fig. 2)

Table S9. (*Excel file*) LC-MS/MS identification of co-purified proteins with pre-CrRbcS2-FLAG_3x_:Protein A:HIS_6x_ translocation intermediates.

(Supplemental data for Fig. 4 and Table 2)

Table S10. (*Excel file*) LC-MS/MS identification of proteins upon depletion of Ti214 in Chlamydomonas cells.

(Supplemental data for Fig. 7)

Table S11. Chlamydomonas proteins retaining their chloroplast transit peptide upon depletion of CrTi214

(Supplemental data for Fig. 7)

**Table S12.** Putative Arabidopsis orthologs of chloroplast protein precursors detected upon depletion of Tic214 in Chlamydomonas.

| Chlamydomonas |  | Arabidopsis |  |  |
| --- | --- | --- | --- | --- |
| Gene ID | Gene Name | Ortholog Gene ID | Ortholog Gene Name | Ortholog Relationship |
| Cre03.g144707 |  | At1g70820 |  | one-to-one |
| Cre03.g146187 |  | At2g19940 |  | one-to-one |
| Cre03.g189800 |  | At3g01480 | *CYP38* | one-to-one |
| Cre06.g273700 |  | At5g23120 | *HCF136* | one-to-one |
| Cre06.g282000 | *STA3* | At1g11720 | *SS3* | one-to-one |
| Cre06.g284750 |  | At1g18070 |  | one-to-one |
| Cre08.g364800 |  | At1g74260 | *PUR4* | one-to-one |
| Cre09.g411200 | *TEF5* | At1g71500 |  | one-to-one |
| Cre10.g433000 |  | At3g48110 | *EDD1* | one-to-one |
| Cre11.g481500 |  | At4g26900 | *AT-HF* | one-to-one |
| Cre12.g497300 | *CAS1* | At5g23060 | *CaS* | one-to-one |
| Cre12.g500650 | *RNB2* | At5g02250 | *EMB2730* | one-to-one |
| Cre16.g663900 | *PBGD1* | At5g08280 | *HEMC* | one-to-one |
| Cre17.g719900 | *PWD1* | At5g26570 | *PWD* | one-to-one |
| Cre48.g761197 |  | At2g43030 |  | one-to-one |
| Cre01.g061077 |  | At1g16880 |  | one-to-many |
| Cre01.g061077 |  | At5g04740 |  | one-to-many |
| Cre02.g080200 | *TRK1* | At2g45290 |  | one-to-many |
| Cre02.g080200 | *TRK1* | At3g60750 |  | one-to-many |
| Cre02.g090850 | *CLPB3* | At2g25140 | *CLPB4* | one-to-many |
| Cre02.g090850 | *CLPB3* | At5g15450 | *CLPB3* | one-to-many |
| Cre03.g158000 |  | At3g48730 | *GSA2* | one-to-many |
| Cre03.g158000 |  | At5g63570 | *GSA1* | one-to-many |
| Cre03.g181300 |  | At1g48860 |  | one-to-many |
| Cre03.g181300 |  | At2g45300 |  | one-to-many |
| Cre05.g234638 |  | At2g16570 | *ASE1* | one-to-many |
| Cre05.g234638 |  | At4g34740 | *ASE2* | one-to-many |
| Cre05.g234638 |  | At4g38880 | *ASE3* | one-to-many |
| Cre06.g250100 | *HSP70B* | At4g24280 | *cpHsc70-1* | one-to-many |
| Cre06.g250100 | *HSP70B* | At5g49910 | *cpHsc70-2* | one-to-many |
| Cre07.g340900 |  | At2g28305 | *LOG1* | one-to-many |
| Cre07.g340900 |  | At2g35990 |  | one-to-many |
| Cre07.g340900 |  | At2g37210 |  | one-to-many |
| Cre07.g340900 |  | At3g53450 |  | one-to-many |
| Cre07.g340900 |  | At4g35190 |  | one-to-many |
| Cre07.g340900 |  | At5g03270 |  | one-to-many |
| Cre07.g340900 |  | At5g06300 |  | one-to-many |
| Cre07.g340900 |  | At5g11950 |  | one-to-many |
| Cre10.g423650 | *PRPL11* | At1g32990 | *PRPL11* | one-to-many |
| Cre10.g423650 | *PRPL11* | At5g51610 |  | one-to-many |
| Cre12.g485800 | *FTSH1* | At1g50250 | *FTSH1* | one-to-many |
| Cre12.g485800 | *FTSH1* | At5g42270 | *VAR1* | one-to-many |
| Cre12.g526800 |  | At1g64190 |  | one-to-many |
| Cre12.g526800 |  | At3g02360 |  | one-to-many |
| Cre12.g526800 |  | At5g41670 |  | one-to-many |
| Cre12.g518900 |  | At2g27680 |  | many-to-one |
| Cre07.g339150 |  | At1g55490 | *CPN60B* | many-to-many |
| Cre07.g339150 |  | At3g13470 |  | many-to-many |
| Cre07.g339150 |  | At5g56500 |  | many-to-many |
| Cre12.g541800 |  | At1g20380 |  | many-to-many |
| Cre12.g541800 |  | At1g76140 |  | many-to-many |

(Supplemental data for Fig. 7)

Table S13. (*Excel file*) List of proteins encoded by Chlamydomonas chloroplast stress-responsive and *MARS1*-dependent genes detected by mass spectrometry depletion of Ti214

(Supplemental data for Fig. S8)

